# Measuring DNA mechanics on the genome scale

**DOI:** 10.1101/2020.08.17.255042

**Authors:** Aakash Basu, Dmitriy G. Bobrovnikov, Zan Qureshi, Tunc Kayikcioglu, Thuy T. M. Ngo, Anand Ranjan, Sebastian Eustermann, Basilio Cieza, Michael T. Morgan, Miroslav Hejna, H. Tomas Rube, Karl-Peter Hopfner, Cynthia Wolberger, Jun S. Song, Taekjip Ha

## Abstract

Mechanical deformations of DNA such as bending are ubiquitous and implicated in diverse cellular functions^1^. However, the lack of high-throughput tools to directly measure the mechanical properties of DNA limits our understanding of whether and how DNA sequences modulate DNA mechanics and associated chromatin transactions genome-wide. We developed an assay called loop-seq to measure the intrinsic cyclizability of DNA – a proxy for DNA bendability – in high throughput. We measured the intrinsic cyclizabilities of 270,806 50 bp DNA fragments that span the entire length of *S. cerevisiae* chromosome V and other genomic regions, and also include random sequences. We discovered sequence-encoded regions of unusually low bendability upstream of Transcription Start Sites (TSSs). These regions disfavor the sharp DNA bending required for nucleosome formation and are co-centric with known Nucleosome Depleted Regions (NDRs). We show biochemically that low bendability of linker DNA located about 40 bp away from a nucleosome edge inhibits nucleosome sliding into the linker by the chromatin remodeler INO80. The observation explains how INO80 can create promoter-proximal nucleosomal arrays in the absence of any other factors^2^ by reading the DNA mechanical landscape. We show that chromosome wide, nucleosomes are characterized by high DNA bendability near dyads and low bendability near the linkers. This contrast increases for nucleosomes deeper into gene bodies, suggesting that DNA mechanics plays a previously unappreciated role in organizing nucleosomes far from the TSS, where nucleosome remodelers predominate. Importantly, random substitution of synonymous codons does not preserve this contrast, suggesting that the evolution of codon choice has been impacted by selective pressure to preserve sequence-encoded mechanical modulations along genes. We also provide evidence that transcription through the TSS-proximal nucleosomes is impacted by local DNA mechanics. Overall, this first genome-scale map of DNA mechanics hints at a ‘mechanical code’ with broad functional implications.

## Measuring DNA mechanics in high throughput

DNA looping (or cyclization) assays have long been used to measure DNA bendability^3,4^. Recently, a single molecule Fluorescence Resonance Energy Transfer^5^ (smFRET)-based DNA looping assay was developed^6^, whereby looping of a ~100 basepair (bp) DNA duplex flanked by complementary 10 nucleotide (nt) single-stranded overhangs is detected via an increase in FRET between fluorophores located at each end of DNA (Fig. 1a). The looping rate thus obtained has been interpreted as a measure of DNA bendability. In this assay, chemically synthesized single strands of DNA had to be annealed directly without PCR amplification to generate a duplex region flanked by long 10 nt overhangs. We simplified the process by developing a nicking-based method that allows the *in situ* conversion of a 120 bp duplex DNA (which can be produced via PCR amplification) into a 100 bp duplex flanked by 10 nt single-stranded overhangs (Fig. 1b). Using FRET^6^, we measured the looping times of ten DNA fragments with different sequences (Supplementary Note 1). The looping times varied by more than an order of magnitude (Fig. 1c), confirming the previously reported result that DNA sequence can have a profound effect on DNA looping at the ~100 bp length scale^6,7^. However, looping assays and all previous methods to directly measure DNA bendability have limited throughput, which greatly limits our knowledge of how DNA mechanics is modulated by sequence, varies along genomes, and influences chromosome transactions.

**Figure 1:**
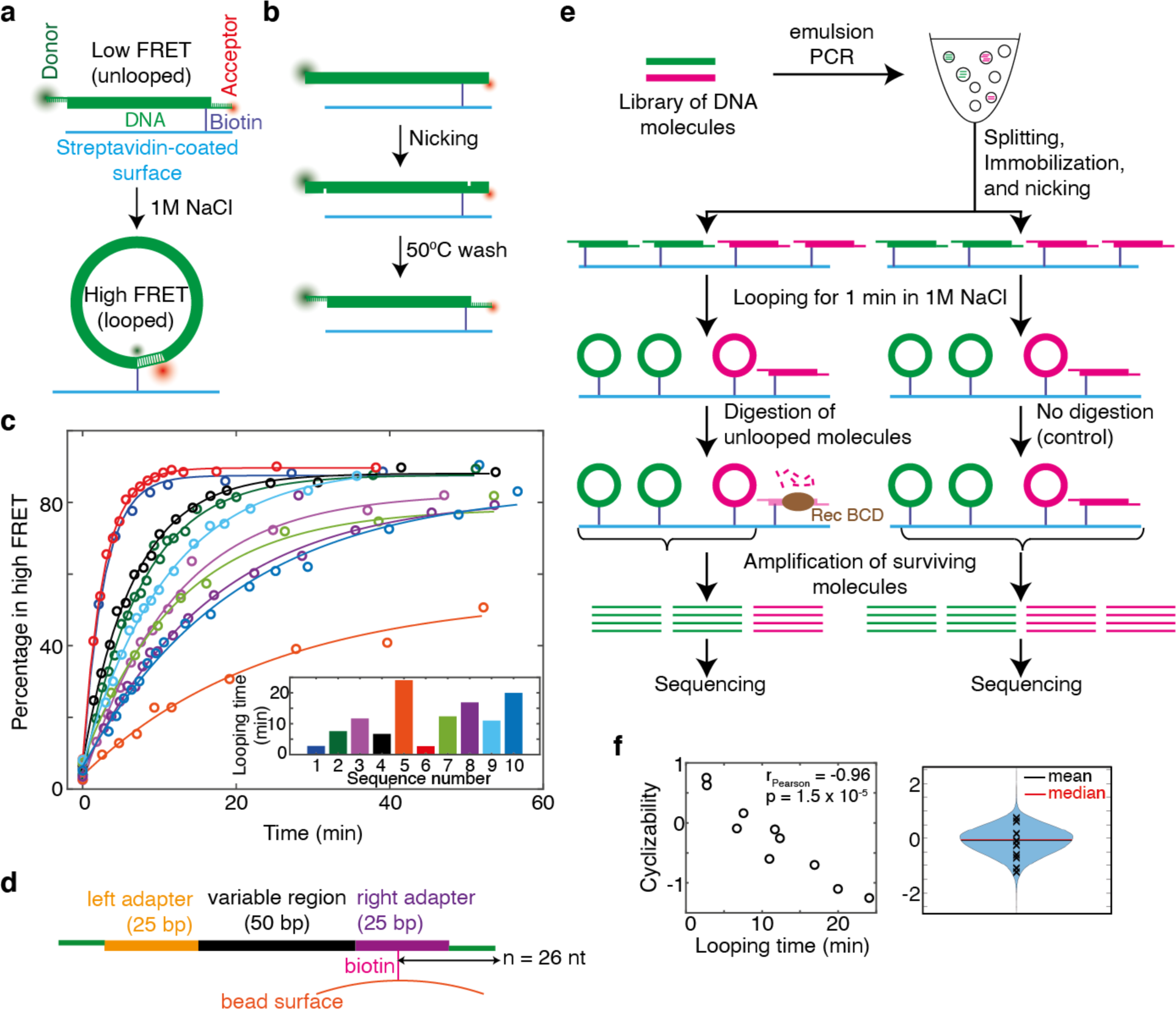
A high-throughout method to measure the mechanical properties of DNA. **(a)** Schematic of the single-molecule looping assay, showing how looping results in an increase in FRET. **(b)** *In situ* nicking of 120 bp duplex DNA 10 nt from either end, following by washing with buffer at 50 °C, results in 100 bp duplex molecules flanked by 10 nt single-stranded overhangs. Site-specific nicking is achieved by engineered recognition sequnces of the nicking enzyme Nt.BspQ1 (NEB). See Supplementary Note 1. **(c)** Percentage of DNA molecules in the high FRET (looped) state as a function of time after adding high salt, as observed for 10 different DNA sequences (Supplementary Note 1). Only molecules that contained both the FRET donor and acceptor were considered. Solid lines represent exponential decay fits, whose characteristic time constant is defined as the looping time and is listed for the 10 sequences in the bar plot (Supplementary Note 1). **(d)** Schematic of a typical DNA fragment in a library just prior to looping. ‘*n*’ is the distance in nucleotides of the biotin tether from the end of the molecule **(e)** Schematic of loop-seq performed on a hypothetical library comprising only two sequences – green and pink. The library is amplified via emulsion PCR^35^, which prevents improper annealing among different library members via the common adapter sequences at the ends. The amplified library is immobilized on beads via biotin-streptavidin interactions and nicked *in situ* to generate loopable molecules (Fig. 1d). After looping for 1 minute in high salt followed by the digestion of unlooped molecules and the amplification of surviving molecules, the relative populations of green and pink in the digested fraction are 2/3 and 1/3 respectively. The corresponding values in the control fraction are 1/2 and 1/2. The cyclizability of green is thus 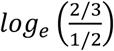 and that of pink is 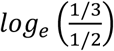. **(f)** Left panel: Cyclizabilities of 10 sequences (listed in Supplementary Note 1) which were part of the Cerevisiae Nucleosomal Library (Supplementary Note 4) as obtained by loop-seq vs looping times of those 10 sequences obtained from smFRET experiments (Fig. 1c). Right panel: Violin plot of the cyclizabilities of all 19,907 sequences in the Cerevisiae Nucleosomal Library. Cyclizabilities of the 10 sequences whose looping times were measured via smFRET are indicated by ‘x’.

Systematic Enrichment of Ligands by Exponential Enrichment (SELEX) has been used to enrich DNA sequences that are more bent^8^ or more loopable^9^ through many rounds of selection of rapidly looping DNA from a vast random pool of loopable molecules and PCR amplification of selected molecules. These assays revealed, for example, certain periodic dinucleotide distributions found in the variable regions of highly loopable DNA. However, direct bendability measurements of specified sequences of interest, such as those than span genomic regions, have never been reported in high throughput.

In order to extend direct DNA bendability measurements to a much larger sequence space, we established a sequencing-based approach termed loop-seq, which builds on previous low-throughput single-molecule looping^6^ and SELEX selection methods^9^. Using the nicking approach, we generated a library of up to ~90,000 different specified template sequences immobilized on streptavidin-coated beads. Library members had a central 50 bp duplex region of variable sequence flanked by 25 bp left and right duplex adapters and 10 nt single-stranded complementary overhangs (Fig. 1d). Looping was initiated in high salt for 1 minute, after which unlooped DNA molecules were digested with an exonuclease^9^ (RecBCD) that requires free DNA ends, thus preserving the looped molecules. The enriched library was sequenced, and the cyclizability of each sequence was defined as the natural logarithm of the ratio of the relative population of that sequence in the enriched library to that in an identically treated control in which only the digestion step was omitted (Fig. 1e, Supplementary Note 2).

The looping times of the 10 sequences determined via smFRET (Fig. 1c) were strongly anti-correlated with their cyclizability values obtained by performing loop-seq on a large library containing those 10 (along with 19,897 other) sequences (Fig. 1f). This confirmed that cyclizability is a good measure of looping rate. Additionally, varying the time for which looping is permitted before RecBCD digestion allowed for the measurement of the full looping kinetic curves of all sequences in the library (‘Timecourse loop-seq’, Supplementary Note 3 and Extended Data Fig. 2). The looped population could comprise closed structures with alternate shapes and basepairing geometries^10,11^ (Extended Data Fig. 1). However, irrespective of looped geometry, control experiments indicate that most looped molecules are protected from RecBCD digestion, and also serve to validate several other aspects of the assay (Extended Data Fig. 3).

We found that the distance of the biotin tether from the end of each molecule (*n*, Fig. 1d) imposed an oscillatory modulation on cyclizability, possibly owing to a sequence-dependent preference for the rotational orientation of the biotin tether (Supplementary Note 7). By varying *n* and performing loop-seq multiple times, we measured the mean, amplitude, and phase associated with this oscillation for every library sequence. We called the mean term the “intrinsic cyclizability” and showed that it is independent of the tethering geometry and rotational phasing (Supplementary Notes 7-8, Extended Data Fig. 4). Both dynamic flexibility and static bending may contribute to intrinsic cyclizability. Regardless of interpretation, intrinsic cyclizability is a measurable mechanical property that can be compared to functional properties of chromosomal DNA.

## DNA at yeast NDRs is rigid

We used loop-seq to query the role of DNA mechanical properties in establishing characteristic features of genes that regulate expression, such as Nucleosome Depleted Regions (NDRs) upstream of the Transcription Start Sites (TSSs) and well-ordered arrays of downstream nucleosomes positioned at characteristic distances from the TSSs^12^. Although several lines of evidence had suggested that DNA mechanics, in addition to transcription factors and chromatin remodelers^2,13,14^, plays a role in this regard by modulating nucleosome organization^2,9,15,16^, the mechanical properties of DNA along promoters and genes have never been directly measured. We measured the intrinsic cyclizabilities of DNA fragments (‘Tiling Library’, Supplementary Note 9) that tile the region from 600 bp upstream to 400 bp downstream of the +1 nucleosome dyads of 576 genes in *S. cerevisiae* at 7 bp resolution (Fig. 2a). We discovered a sharply defined region of rigid DNA (i.e. with unusually low intrinsic cyclizability) located in the NDR^17^ (Fig. 2b). Further, there are many genes where our measurements are sensitive enough to detect this region of high rigidity without the need to average across multiple genes (Fig. 2c, Extended Data Fig. 5). As nucleosome assembly requires extensive DNA bending, the low intrinsic cyclizability of DNA around the NDR is likely to favor nucleosome depletion.

**Figure 2:**
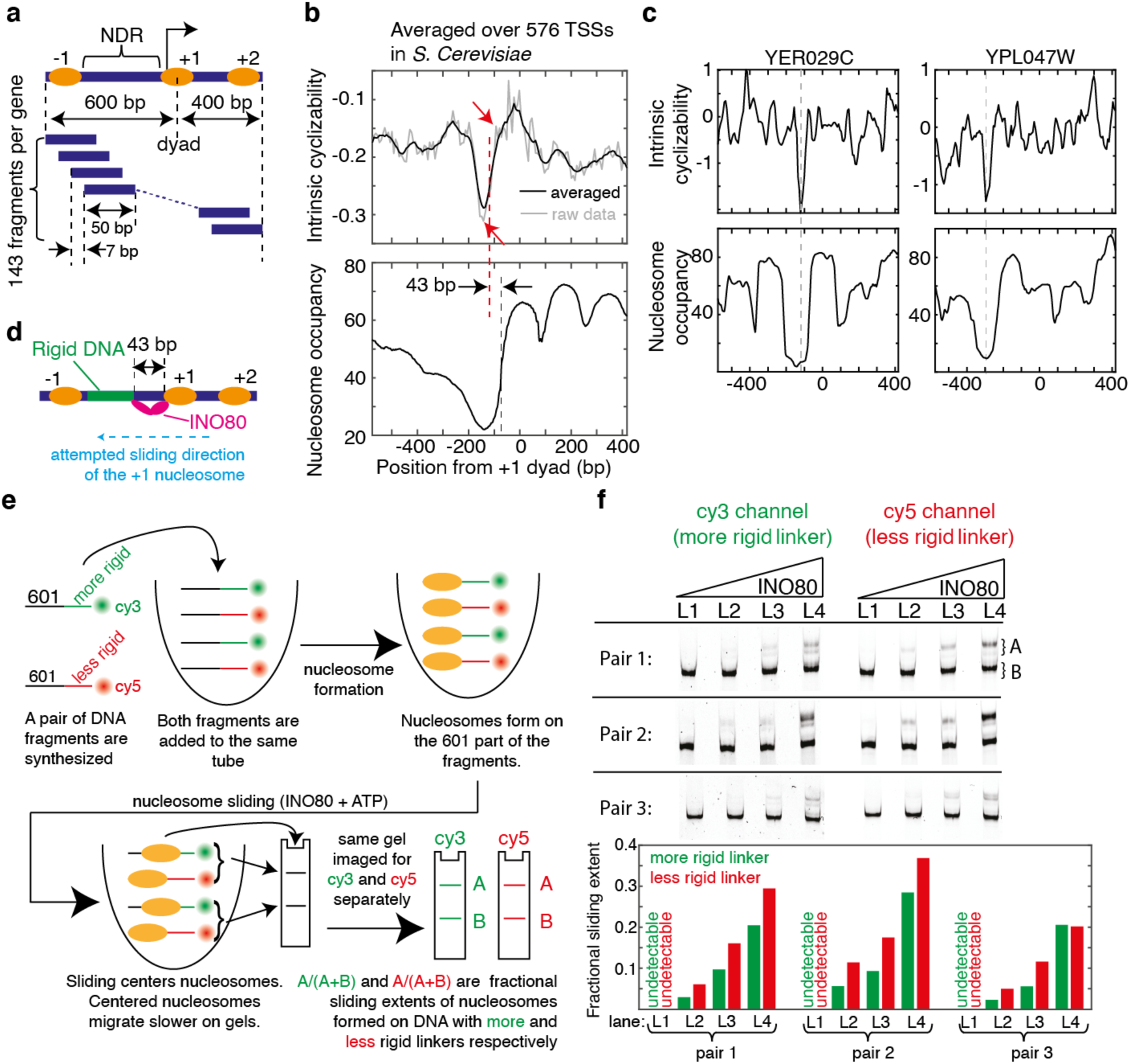
DNA mechanics contributes to nucleosome depletion at the NDR and modulates remodeler activities. **(a)** Schematic of the Tiling Library. The central 50 bp variable region was designed to tile 576 genes in *S. cerevisiae* at 7 bp resolution from 600 bp upstream till 400 bp downstream of the dyads of the +1 nucleosomes of these genes every 7 bp (Supplementary Note 9). DNA is represented by the blue lines, and nucleosomes by the oval shapes. **(b)** Mean intrinsic cyclizability (with (black) and without (grey) any smoothening) and nucleosome occupancy vs position from the canonical location of the dyad of the +1 nucleosome, averaged over all 576 genes in the Tiling Library. Nucleosome occupancy values^17^ and positions^32^ were as reported earlier. All genes are oriented in the direction of transcription. The blue dashed line (at −73 bp) marks the upstream edge of the +1 nucleosome. The red dashed line marks the start of the rigid DNA region, approximated as the midpoint between the two red arrows. See supplementary note 10. **(c)** Intrinsic cyclizability and nucleosome occupancy vs position from the dyads of the +1 nucleosomes of two individual genes (see Extended Data Fig. 5 for six more examples). **(d)** Schematic showing an INO80 enzyme attempting to slide a +1 nucleosome upstream of its canonical location. The rigid DNA region starts ~40 bp upstream of the +1 nucleosome edge (Fig. 2b), about where the Arp8 module of an INO80 attempting to slide the nucleosome upstream would bind ^21,23^. **(e)** Schematic of the experiment comparing the effect of DNA rigidity on nucleosome sliding by INO80. A pair of constructs were engineered involving a nucleosome formed on the 601 DNA sequence attached to two different 80 bp linker DNAs that had a large difference in intrinsic cyclizability values around the middle of the 80 bp region. The construct with the lower intrinsic cyclizability (i.e. more rigid) 80 bp region was labeled with Cy3 at the 5’ terminus distal to the 601 region and the other construct was labeled with Cy5. Both constructs were present in the same reaction volume, and sliding was effected for 1 minute in the presence of INO80 and ATP. The products were run on a gel. Sliding results in the generation of centered nucleosomes, which migrate more slowly^23^. The gel was imaged separately for Cy3 and Cy5 fluorescence to quantify the extent of sliding of nucleosomes formed on the two constructs in the pair independently. See Supplementary Note 11. **(f)** Three separate nucleosome sliding experiments were performed involving three different pairs of nucleosome constructs as described in figure 2e and supplementary note 11. In each experiment, four concentrations of INO80 were used (in lanes L1 – L4): 2, 6, 9, and 13 nM. The products of the sliding reaction were run along a 6% TBE gel, which was imaged separately for Cy3 and Cy5 fluorescence. For each concentration of INO80, i.e. for each lane, the nucleosome sliding extents in the two constructs in the pair were quantified (bar plots) as described in figure 2e and supplementary note 11.

## Chromatin remodelers sense DNA mechanics

Chromatin remodelers have been proposed to be critical in establishing the well-ordered array of nucleosomes downstream of TSSs by stacking nucleosomes against a barrier just upstream of the TSS^14^. What could constitute such a barrier has been a matter of debate, and transcription factors^13^ and paused polymerases^18^ have been suggested to contribute. Notably, *in vitro* chromatin reconstitution experiments^2^ showed that the remodeler INO80 can both position the +1 (and −1) nucleosomes and establish the NDRs in *S. cerevisiae* even in the absence of any such factors. We therefore asked whether the sequence-encoded rigid DNA region in the NDR can contribute to nucleosome positioning near promoters by serving as a barrier to the sliding activities of INO80.

To effect sliding, INO80 requires at least 40 – 50 bp of free extranucleosomal DNA ahead of the nucleosome^19,20^. The region around 40 – 50 bp ahead of the sliding nucleosome’s edge is enganged by the Arp8 module of INO80^21–23^, and disrupting the module’s DNA binding via mutation abolishes sliding and reduces +1 positioning genome-wide^23^. Intriguingly, we found that the region of rigid DNA also starts ~43 bp upstream of the edge of the +1 nucleosome (Fig. 2b). This would place the Arp8 module in contact with highly rigid DNA if INO80 were to slide the +1 nucleosome upstream from its canonical position (Fig. 2d). If highly rigid DNA interferes with Arp8 module binding, further upstream sliding of the +1 nucleosome would be hindered, helping position the +1 nucleosome and define the NDR.

To directly test the role of upstream DNA rigidity in +1 nucleosome positioning by INO80, we biochemically measured the effect of rigid DNA located ~40 bp ahead of a nucleosome on sliding by INO80. Using gel shift, we assayed sliding of nucleosomes formed on the 147 bp 601 sequence into adjacent 80 bp linkers. We chose three pairs of such constructs, each containing one construct with a linker that was uniformly flexible and another with a linker that had a significantly more rigid region near the middle (Supplementary Note 11, Extended Data Fig. 6a). In all three pairs, the extent of sliding (Supplementary Note 11) was lower for the nucleosome formed on the construct with the rigid linker (Fig. 2f, Extended Data Figs. 6b-c, 7). Various factors could cause this reduced sliding (Supplementary Note 11). Regardless, the observation is consistent with a model where the rigid DNA region starting ~43 bp upstream of the canonical +1 nucleosome’s edge (Fig. 2b) serves as a barrier that hinders further upstream sliding of the +1 nucleosome by INO80, possibly aided by other barriers set up by factors such as RSC, gene regulatory factors, and transcription factors^2,13^. Structural details behind rigidity sensing by the Arp8 module must await future studies.

## DNA mechanics impacts nucleosome organization

As nucleosomes involve extensive DNA bending, we asked if modulations in intrinsic cyclizability may directly contribute to nucleosome organization, in addition to the stacking action of remodelers^14^. Indeed, DNA at the canonical dyad locations of the +/-1 nucleosomes, and to a lesser extent the +2, +3 and +4 nucleosomes, have significantly higher intrinsic cyclizability than surrounding DNA (Fig. 2b). Consistent with this observation, promoters classified as having a fragile −1 nucleosome^24^ have more rigid DNA at the location of the −1 nucleosome (Fig. 3a).

**Figure 3.**
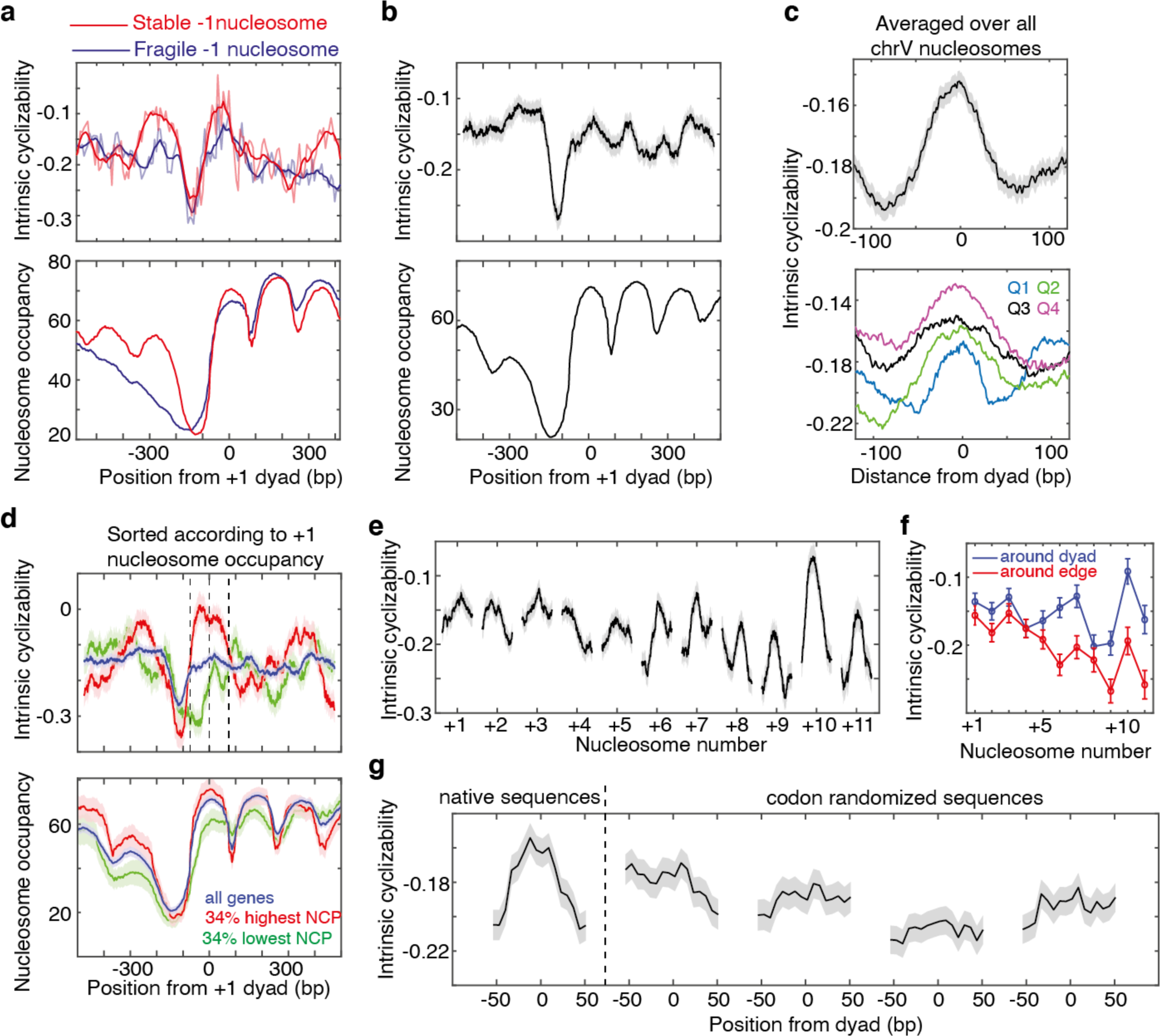
DNA mechanics contributes to global nucleosome organization. **(a)** Mean intrinsic cyclizability and nucleosome occupancy vs position from the dyad of the +1 nucleosome, averaged separately over 185 and 345 genes in the tiling library that have been classified as possessing stable and fragile −1 nucleosomes respectively^24^. Plots were generated as in figure 2b. **(b)** Mean intrinsic cyclizability and nucleosome occupancy vs position from the +1 nucleosomal dyad, averaged over 227 identified genes along *S. cerevisiae* chromosome V. Grey background represents s.e.m. See supplementary note 13. **(c)** Top: intrinsic cyclizability vs position from the dyad, averaged over all 3,192 nucleosomes in chromosome V. Shaded background represents s.e.m. Bottom: Nucleosomes were sorted into quartiles based on reported NCP scores^32^, and average flexibility profiles across nucleosomes were plotted for each quartile. See Extended Data Fig. 8a-b for quartile boundaries, and for plots along octiles. **(d)** Intrinsic cyclizability and nucleosome occupancy vs position along all genes in chromosome V (blue) and among the 34% of these genes that had the highest (red) and lowest (green) +1 nucleosome NCP scores^32^. Plots were obtained as in panel b. The three vertical dashed lines mark the edges and dyad of the +1 nucleosome. Shaded backgrounds represent s.e.m. **(e)** Mean (solid line) and s.e.m. (grey background) of intrinsic cyclizability values around the dyads of gene body nucleosomes that lie between the TSS and the TTS of all identified 227 genes along chromosome V in *S. cerevisiae*. **(f)** Mean and s.e.m. of intrinsic cyclizability values of DNA in a 50 bp window around the dyads of gene body nucleosomes (blue) and in a 50 bp window around the edges of gene body nucleosomes (from positions −73 till −56 and from +56 till +73 with respect to the dyad) (red). Solid lines are guides to the eye. **(g)** Left of dashed line: intrinsic cyclizability as a function of position along the native sequences of the 500 +7 nucleosomes represented in Library L (supplementary note 14). Right of dashed line: same quantity for the four sets of codon-altered sequences. The first two plots are for the cases where synonymous codons were selected considering the natural codon-usage frequency in *S. Cerevisiae*. This was not the case for the subsequent two plots. Solid line represents mean, and the grey background represents s.e.m. Data were plotted with a rolling window averaging of 7 adjacent fragments.

Several earlier studies have shed light on the role of DNA mechanics in nucleosome formation^25^. The fact that bendable DNA forms good substrates for nucleosomes and vice versa, has been demonstrated for various selected sequences^4,26–30^. Further, DNA selected for high loopability from a large random pool possess periodic distribution in dinucleotide contents^9^, which is also a feature found in ~3% of native yeast nucleosomal sequences^31^. However, the mechanical properties of known nucleosomal DNA sequences have never been directly measured in high throughput. To achieve this for nucleosomes along an entire yeast chromosome, we measured intrinsic cyclizability along *S. cerevisiae* chromosome V at 7 bp resolution (‘ChrV Library’, Supplementary Note 12, Extended Data Fig. 8). We first confirmed that intrinsic cyclizability shows the characteristic pronounced dip around the NDR when averaged over the 227 genes along chromosome V that have both ends mapped with high confidence^25^ (Fig. 3b). We found that chromosome-wide, DNA at nucleosomal dyad locations tends to have significantly higher intrinsic cyclizability than the surrounding linker DNA (Fig. 3c), suggesting that sequence-dependent modulations in DNA mechanics contribute to global nucleosome organization. We also found that nucleosomes are better positioned *in vivo* on more intrinsically cyclizable DNA (Fig. 3c, Extended Data Fig. 9a-c). Among TSS proximal nucleosomes, the correlation is strongest for +1 nucleosomes (Fig. 3d, Extended Data Fig. 9d-e).

Among nucleosomes that lie along transcribed regions, TSS-distal nucleosomes have a higher intrinsic cyclizability contrast between the dyad and the edges than TSS-proximal nucleosomes (Figs. 3e–f). This was contrary to expectation because TSS-proximal nucleosomes are known to be better positioned than TSS-distal nucleosomes^14,32^. TSS-proximal nucleosomes are likely primarily organized by chromatin remodelers into ordered arrays via stacking against the NDR barrier^2,14^. However, beyond the +4 nucleosome, the stacking effect has been shown to dissipate^14^, whereas our data shows that modulations in intrinsic cyclizability become more prominent (Fig. 3e–f). Thus nucleosomes that lie deeper in gene-bodies may rely more on sequence-encoded intrinsic cyclizability modulations for positioning.

## Codon choice optimizes mechanical modulations

We next asked whether the strong modulation in intrinsic cyclizability for nucleosomes deep in the gene body would be preserved if the sequences were altered by using alternate codons that code for the same amino acids. We selected 500 +7 nucleosomes in *S. cerevisiae* and generated four sets of codon-altered sequences spanning the region around these nucleosomes, while preserving the amino acid sequences encoded. The natural codon usage frequency was considered when choosing synonymous codons in the first two sets, and was ignored in the next two (supplementary note 14). By performing loop-seq (‘Library L’, supplementary note 14), we measured intrinsic cyclizability at 7 bp resolution in the 200 bp region flanking the 500 +7 nucleosome dyads and their codon-altered sequences. Native sequences have a characteristic intrinsic cyclizability pattern – high near the dyads and low near the edges – which is absent in the four codon-altered sets (Fig. 3g). Thus naturally occurring codons are optimized to establish sequence-dependent intrinsic cyclizability modulations along genes that are favorable to the organization of gene body nucleosomes, suggesting that the evolution of codon choice in *S. cerevisiae* has been impacted by a selective pressure to preserve such modulations. The observation also points to a hitherto unappreciated significance of positioning nucleosomes that lie deeper in the gene body.

## TSS-proximal nucleosomes are asymmetric

Several critical processes such as transcription and DNA replication require nucleosome unraveling. DNA could potentially peel off from either end, in a manner modulated by bendability. Indeed asymmetry in DNA bendability across the 601 nucleosome leads to asymmetric unraveling under tension^33^. Biochemical analysis has shown that yeast RNA polymerase II transcribing a 601 nucleosome produces four times more full-length transcripts when it enters the nucleosome through the ‘TA-rich’ side that contains the phased TA repeats^34^. Using loop-seq, we also found that the TA-rich side has significantly higher intrinsic cyclizability (Fig. 4a, Supplementary Note 18). This observation is consistent with an idea that RNA polymerase might better negotiate with a nucleosomal barrier when it first interacts with the side of the nucleosome containing DNA with higher intrinsic cyclizability. We constructed a library containing the 50 bp DNA fragments immediately to the left and right of the dyads of ~10,000 well-positioned *S. cerevisiae* nucleosomes (‘Cerevisiae Nucleosomal Library’, Supplementary Note 4, Fig. 4b). We found that DNA at well-occupied +1 and +2 nucleosomes indeed has, on average, higher intrinsic cyclizability on the promoter-proximal face than the distal face (Fig. 4c), thereby suggesting that this asymmetry may favor polymerase translocation. Consistently, this asymmetry is accentuated among the highly expressed genes and absent among poorly expressed genes (Fig. 4d).

**Figure 4:**
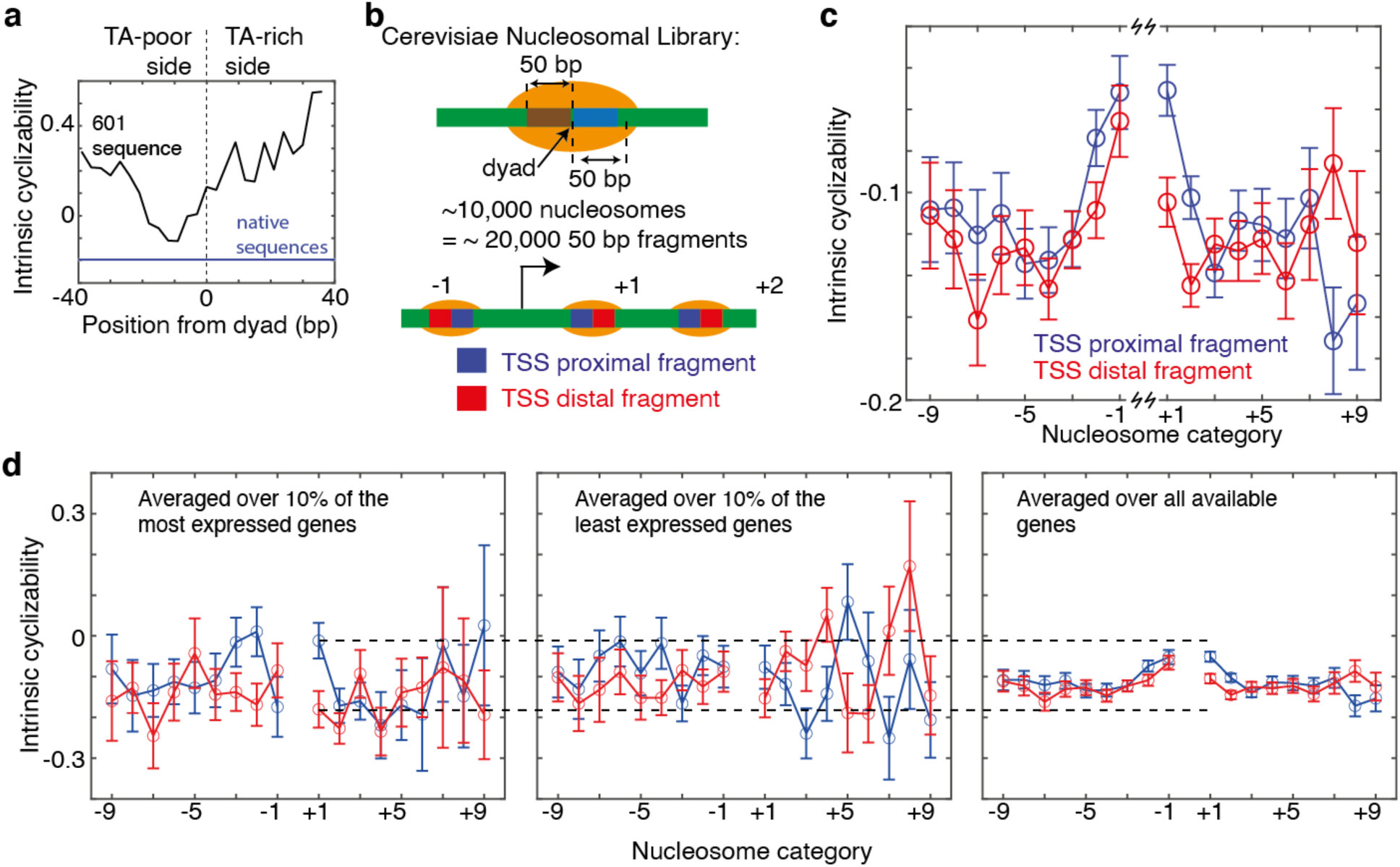
TSS-proximal nucleosomes are asymmetric. **(a)** Black: Intrinsic cyclizability as a function of position along 601 DNA (supplementary note 18) Blue: abscissa value of the solid horizontal line (−0.196) is the mean intrinsic cyclizability along the 500 native +7 nucleosomal sequences represented in Library L (supplementary note 14). The height of the light blue background (0.011) is twice the s.e.m. **(b)** Schematic representing the design of the Cerevisiae Nucleosomal Library (supplementary note 4). The library contains DNA fragments taken from the 50 bp immediately to the left and right of the dyads of the ~10,000 nucleosomes in *S. cerevisiae* that have the highest NCP scores. **(c)** Mean intrinsic cyclizabilities of the 50 bp DNA fragments that lie immediately adjacent to the TSS proximal (red) or distal (blue) side of the dyads of various categories of nucleosomes (−9 through +9) (see supplementary note 17). Error bars are s.e.m. **(d)** A subset of the data as in panel c, where the means were calculated considering only genes among the 10% most (left panel) or least (middle panel) expressed in *S. cerevisiae*. The right panel is identical to panel b, except for an altered y-axis scale. Error bars are s.e.m. See supplementary note 17.

## Conclusions

Intrinsic cyclizability is, thus far, the only mechanical property of DNA to be directly measured in high throughput, and will likely aid our understanding of how DNA mechanics influences chromatin transactions involving diverse factors such as topoisomerases, transcription factors, polymerases, structure maintenance of chromatin proteins and so on. The large dataset enabled by loop-seq should make it possible to develop comprehensive models to predict intrinsic cyclizability and other physical properties from DNA sequence. Preliminary analysis showed that simple sequence features such as GC content, polyA tracts, and dinucleotide parameters are generally poor or insufficient predictors of intrinsic cyclizability (Supplementary Note 16, Extended Data Fig. 10).

Our measurements suggest that intrinsic cyclizability is functionally important and must have applied selective pressure throughout the evolution of genomes. It remains to be investigated how genetic information content and the mechanical properties of DNA are linked, and how the sequence-dependent mechanical response of DNA to molecular-scale forces in its immediate environment may have influenced both the slow divergence of organisms and rapid mutations in contexts such as cancer.

## Methods

### smFRET based single-molecule DNA looping assay

Templates were purchased (IDT DNA) and converted into loopable molecules with 10 bp complementary overhangs on either side, Cy3 and Cy5 fluorophores at the ends, and a biotin molecule (Supplementary Note 1) via PCR amplification with KAPA Hi Fi Polymerase (Roche) and nicking near the ends by the site-specific nicking enzyme Nt.BspQ1 (NEB). Molecules were immobilized on a PEG-coated quartz surface (JHU slide production core for microscopy) functionalized with a small amount of biotin-PEG, via a streptavidin sandwich, as described previously^6^. Immobilized molecules were incubated with T2.5 (2.5 mM NaCl, 10 mM Tris-HCl pH 8) for 1.5 hours. Low salt imaging buffer (20 mM Tris-HCl pH 8, 3 mM Trolx, 0.8% dextrose, 0.1 mg/ml glucose oxidase, 0.02 mg/ml catalase) was flowed into the channel and the molecules were imaged on a TIRF microscope to determine the initial histogram of FRET values. High salt imaging buffer (1 M NaCl, and all components of the low salt imaging buffer) was then introduced into the channel at time 0, and FRET histograms were measured at various time points as done previously^6^. The plot of the percentage of molecules with both donor-acceptor pairs in high FRET as a function of time was fit to an exponential. Its time constant was defined as the looping time. The inverse of this was defined to be the looping rate.

### Loop-seq

Instead of individual templates, entire libraries representing as many as ~90,000 individual DNA sequences, with the central 50 bp variable and flanked by identifcal 25 bp adapters, were obtained (Genscript), and amplified using KAPA Hi Fi polymerase (Roche) in 20 cycles of emulsion PCR^35^ (ePCR) using the Micellula DNA emulsion and purification kit (CHIMERx). The manufacturer’s guidelines were followed during ePCR. ePCR prevents improper annealing among different template molecules via the common adapter sequences. Amplification converted the library into 120 bp duplex molecules with a biotin near one end, and the recognition sequence for the nicking enzyme Nt.BspQ1 (NEB) near both ends (Supplementary Note 1). 20 μl of streptavidin-coated magnetic beads (Dynabeads MyOne Streptavidin T1, Thermo Fisher Scientific) were washed 2x with 400 μl T50 BSA (1 mg/ml BSA (Invitrogen) in T50 (50 mM NaCl, 10 mM Tris-HCl pH 8.0)) and resuspended in 20 μl T50 BSA. 2 μl of ~4 ng/ul amplified DNA was mixed with 5 μl of water, and 20 μl of the washed magnetic beads were added. After incubation for 10 minutes, the DNA bound beads were washed 2x with 200 μl T50BSA and 1x with 200 μl T10BSA (1 mg/ml BSA (Invitrogen) in T10 (10 mM Tris-HCl pH 8.0, 10 mM NaCl)). Digestion mix (84 μl water, 10 μl 10x NEB Buffer 3.1, 6 μl Nt.BspQ1 (NEB)) was prepared and heated to 50 oC for 5 minutes. Digestion resulted in an immobilized library, where every DNA molecule has a central 50 bp duplex variable region, flanked by 25 bp left and right adapters and 10 nt complementary single-stranded overhangs (Fig. 1d, Supplementary Note 1). The beads were pulled down and incubated with the heated digestion mix for 25 mins at 37 oC. The beads were then washed 2x with 100 μl of T10BSA preheated to 50 oC, followed by 200 μl of T2.5BSA (1 mg/ml BSA (Invitrogen) in T2.5 (10 mM Tris-HCl pH 8.0, 2.5 mM NaCl)). The beads were incubated in 200 μl T2.5BSA for 1.5 hours on a rotor at room temperature. The bead sample was then split into two 95 μl fractions denoted ‘sample’ and ‘control’. The beads in the sample fraction were pulled down and resuspended in 200 μl looping buffer (1M NaCl, 1 mg/ml BSA, 10 mM Tris-HCl pH 8) for 40 seconds. High salt (1M NaCl) initiates looping, which allows the complementary single-stranded overhangs at the ends to stably hybridize^6^. Apparent DNA bendability has been shown to be independent of the salt concentration used^6^.The tube containing the sample was then placed on magnets for an additional 35 seconds. The looping buffer was replaced with 200 μl of digestion buffer (6.66 μl of RecBCD (NEB), 20 μl NEB 10X Buffer 4, 20 μl of 10x ATP (NEB), 154 ul water) for 20 minutes. This was defined as looping for 1 minute. In general, looping for n minutes implies incubation in looping buffer for up to 20 seconds prior to the completion of n minutes, followed by 35 seconds over magnets before the solution was replaced with digestion buffer. After 20 minutes, digestion buffer was removed by pulling down the beads and replaced with 200 μl of looping buffer. The control was treated in exactly the same way, except the digestion buffer had 6.66 μl of water instead of RecBCD. Beads in the sample and control fractions were then pulled down and the looping buffer was replaced with 50 μl of PCR mix (25 μl 2x HiFi KAPA Hot Start ready mix (Roche), 1 ul each of 100 μM primers (supplementary note 1), 23 μl water) and PCR amplified (16 cycles). If the library contained less than 20,000 sequences, the products were sequenced on an Illumina MiSeq machine. For more complex libraries a HiSeq machine was used. Library preparation for sequencing was done using the Nextera XT primer kit and followed a protocol similar to the Illumina protocol for 16S metagenomic sequencing library preparation.

Sequencing results were mapped to the known sequences in the library using bowtie^36^. The number of times each sequence was represented in the sample and control was obtained and 1 was added to all counts. The relative population of each sequence in the digested and control pools was calculated. Cyclizability of a sequence was defined as the natural logarithm of the ratio of the relative population of a sequence in the sample pool to that in the control. In addition to Bowtie^36^, Samtools^37^ and MATLAB (Matworks) were used to analyze the data.

### Purification of INO80

INO80 was purified according to a protocol published earlier^38^. Briefly, *S. cerevisiae* cells were grown in 12 liters of YPD medium to an O.D. of 1.5. Frozen yeast cells were lysed in a SPEX freezer mill (15 cycles: precool 2 min, run time 1 min, cool time 1 min, rate 15 cps). Ino80-3Flag was affinity-purified from whole lysate using anti-Flag M2 agarose beads and eluted with Flag peptide (0.5 mg/ml). The complex was further purified by sedimentation over a 20-50% glycerol gradient. Peak INO80 fractions were pooled and concentrated using Centricon filters (50 kDa cut off), and buffer changed to 25 mM HEPES–KOH (pH 7.6), 1 mM EDTA, 2 mM MgCl_2_, 10% glycerol, 0.01% NP-40, 0.1 M KCl. Aliquots of purified INO80 were flash-frozen and stored at −80°C. Recombinant INO80 was also purified as per earlier protocols^21^.

### Nucleosome sliding by INO80

Nucleosome preparation and sliding by INO80 was performed under conditions as reported earlier^19^. Sliding for in the presence of various [INO80] as reported in Fig. 2f (for 1 minute) and Extended Data Fig. 7 (for 1 minute and 2.5 minutes) was performed in 10 μl reaction volumes containing 8 nM nucleosomes (nucleosomes formed on both constructs in the pair were present in equimolar proportion), 2 mM ATP, 24 mM tris-HCl pH 7.5, 43 mM KCl, 2.86 mM MgCl2, 0.55% glycerol and indicated concentration of INO80. The mixture was incubated without ATP at 30 oC for 7 minutes. After addition of ATP, the reaction was allowed to proceed for 1 minute at 30 oC, and was then quenched by the addition of lambda DNA and ADP to final concentrations of 66.7 µg/ml and 20 mM respectively. For all sliding experiments reported in Extended Data Fig. 6b (timecourse of INO80 sliding), conditions were the same except incubations prior to ATP addition and the subsequent sliding reaction were carried out at room temperature. The reaction was continued for the indicated amounts of time in presence of saturating INO80 prior to quenching. Quenched reactions were loaded on to 6% TBE gels (Invitrogen) in presence of 10% glycerol and run at 150 V for 1.5 hrs. The gel was imaged separately for Cy3 and Cy5 fluorescence.

## Author Contributions

AB and TH designed research. AB performed research and analyzed data. AB and TH wrote the paper. Other authors contributed in the following areas: DGB – extraction of nucleosome occupancy from published data and plectoneme density calculations. ZQ – preparation of some libraries. TK, TN – initial assay development and the characterization of RecBCD. AR, SE, KPH – purification of INO80 and related insights. BC – observation of the phasing effect. MTM, CW – initial development of nucleosome assays. MH, TR, JSS – initial analysis of sequencing data. All authors commented on the manuscript.

## Data Availability

All raw data and accession codes will be made available upon request.

## Code availability

All codes used to analyze sequencing data will be made available upon reasonable request.

## Acknowledgements

A.B. and T.H. would like to thank Carl Wu, Xinyu Feng, and Matthew F. Poyton for insights and help related to INO80 biochemistry, and Quicen Zhang for help with initial assay development efforts. K.P.H and S.E. would like to thank Manuela Moldt for purification of recombinant INO80. This work was supported by the National Science Foundation Grants PHY-1430124 and EFMA 1933303 (to T.H.), the National Institutes of Health Grants GM122569 (to T.H.), the National Institutes of Health grant NIH R01CA163336 (to J.S.S.), the European Research Council (Advanced Grant INO3D to K.P.H), and by the Deutsche Forschungsgemeinschaft (CRC1064 and Gottfried Wilhelm Leibniz-Prize to K.P.H). A.B. was a Simons Foundation Fellow of the Life Sciences Research Foundation. T.H. is an Investigator with the Howard Hughes Medical Institute.

