## Extended Data Figures 1 - 10 for "Measuring DNA mechanics on the genome scale"

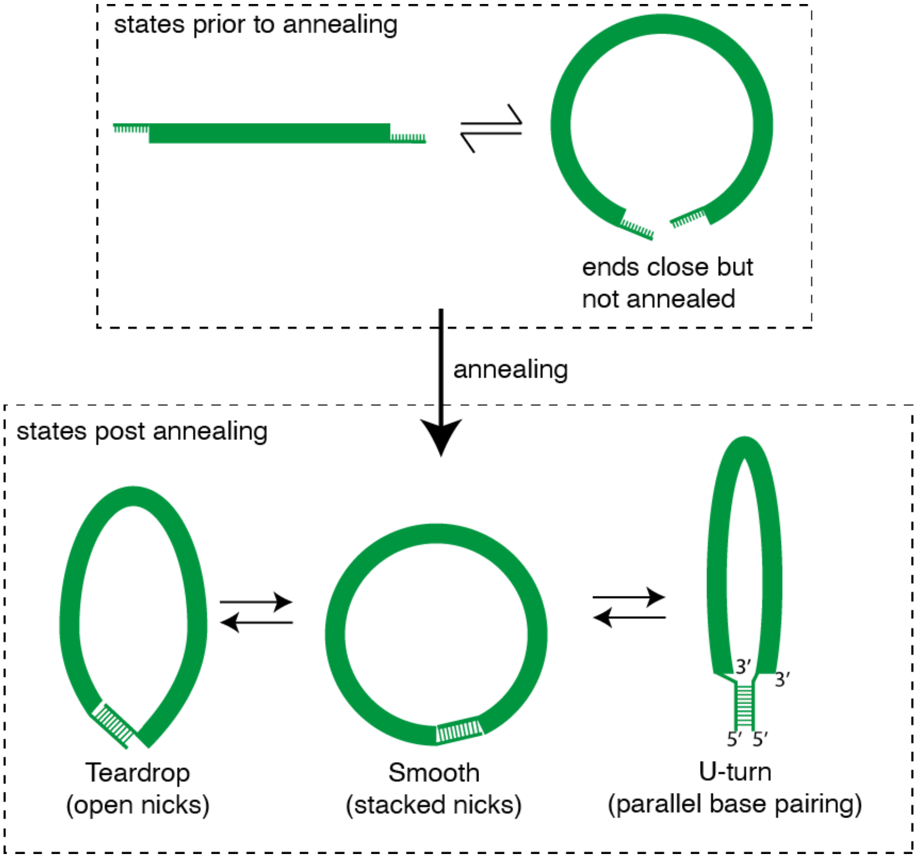


**Extended Data Fig. 1: Pre-looped and looped geometries.** Prior to annealing of the ends, the DNA rapidly samples various configurations where the ends are far apart or closer together, described here for simplicity as a rapid equilibrium between two representative states. As described earlier[1], annealing captures the state where the ends are close together. Thus the rate of looping as measured in the FRET based assay reports on the equilibrium population fraction of the state where the ends are close but not annealed, irrespective of the exact shape and geometry of the subsequent annealed state. It thus addresses the biological question of how quickly regions of DNA can approach, which can then be stabilized by protein binding[1]. However, nucleoprotein complex formation may require not just the ends to approach, but the intervening DNA to also adopt a certain shape, and the readout of looping rate does not distinguish between these possible shapes. Various shapes have been proposed for the subsequent annealed state, such as a teardrop configuration where the nicks are open and basepair stacking across nicks is disrupted, and a smooth state where basepair stacking is preserved across the nicks[2]. Other non-canonical geometries may also be possible, such as a U-turn geometry, where the sticky overhangs interact via reverse Watson-Crick basepairing in a parallel stranded configuration. Conventional Watson-Crick base pairing between the overhangs, but in a geometry similar to the U-turn configuration, has been achieved in the past (hairpin loop[3]). Presence of mismatches or transient defects in the duplex region may influence whether such a geometry is preferred over the teardrop or smooth configuration[3]. Because all members of various libraries in our study used the same overhang sequences, relative differences between them in their looping kinetics are unlikely to be affected by a potential conformational heterogeneity of the looped state.

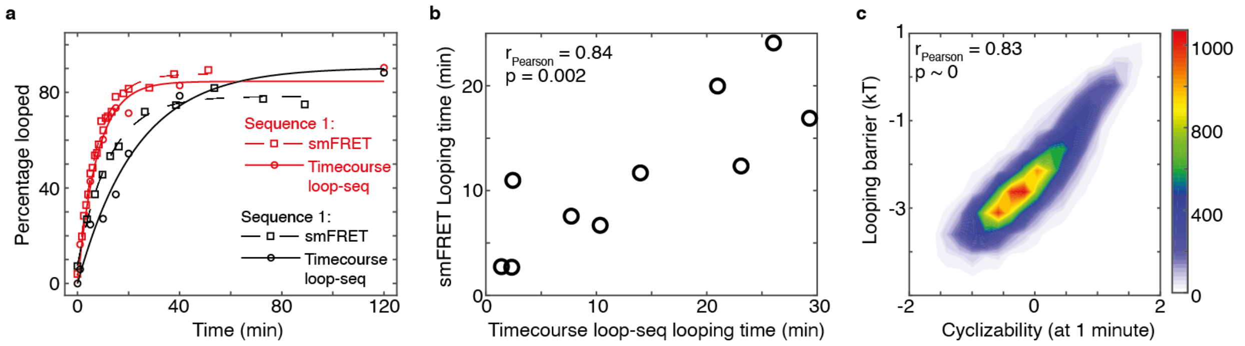


**Extended Data Fig. 2: Timecourse loop-seq. a,** Looping kinetic curves of two individual sequences that were part of the Cerevisiae Nucleosomal Library (Supplementary Note 4), obtained by performing two individual smFRET experiments (Fig. 1a-c) as well as timecourse loop-seq (Supplementary Note 3) on the library. **b,** Looping times of 10 sequences that were part of the Cerevisiae Nucleosomal Library obtained from 10 individual smFRET experiments (Fig. 1c) vs looping times obtained by performing timecourse loop-seq on the library. **c,** Looping barriers (natural logarithm of the looping times, see Supplementary Note 2) of all 19,907 sequences in the Cerevisiae Nucleosomal Library obtained by performing timecourse loop-seq vs the corresponding cyclizability values obtained by performing regular loop-seq involving 1 minute of DNA looping prior to RecBCD digestion.

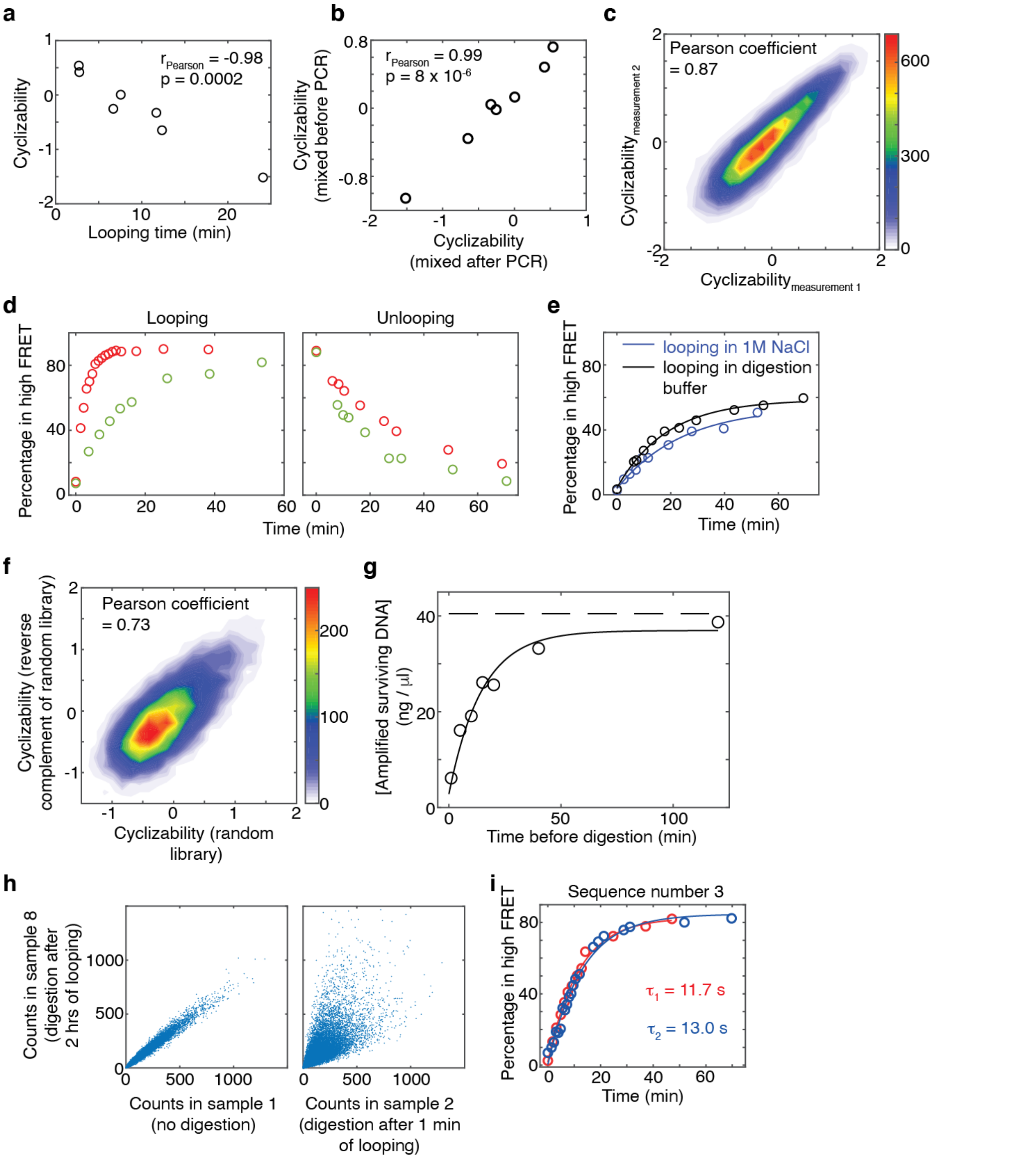


**Extended Data Fig. 3: Controls pertaining to the loop-seq assay. a,** Seven of the ten sequences whose looping times were measured using smFRET (Sequences 1 – 7 in Fig. 1c, and listed in Supplementary Note 1) were combined to form a small library. Plotted are the cyclizability values obtained by performing loop-seq on this library vs the looping times obtained from the individual smFRET experiments. This plot is similar to that in Fig. 1f, except that loop-seq was performed on a much smaller library comprising only these seven sequences. This control serves to revalidate the anti-correlative relation between looping time and cyclizability. **b,** Regular PCR of the entire library containing multiple templates can generate incorrectly annealed products that are annealed via the 25 bp identical adapters at the ends but have 50 bp bubbles in the middle. Such constructs would likely be extremely flexible and would be protected from digestion due to rapid cyclization. Emulsion PCR separates the templates in individual droplets, thereby preventing incorrect annealing between different templates. We performed a control experiment to verify that emulsion PCR of the library does not affect the measured value of cyclizability. In one case, seven template sequences (Sequences 1 – 7 as listed in supplementary note 1 and Fig. 1c) were mixed and then a single round of ePCR was performed to form a small library. In another case, seven separate regular PCR amplifications were carried out for the seven template molecules. The amplified products were then mixed in equimolar proportions to form the library. Loop-seq was performed on these two 7 member libraries and two sets of cyclizabilities of the seven sequences were measured. As indicated by the plot here, these values are highly correlated. **c,** Technical replicates of loop-seq performed on the Cerevisiae Nucleosomal Library (Supplementary Note 4). **d,** This control was performed to verify the expectation that DNA sequences which are more bendable and thus loop quickly under high salt conditions are slow to unloop under low salt conditions. In red is the looping kinetics of Sequence 6, and in green is that of Sequence 7 (Fig. 1c) measured using smFRET. For the unlooping measurements, the slide containing nicked DNA was incubated for 2 hours with high salt imaging buffer containing 1M NaCl. After that, low salt imaging buffer containing no added NaCl was flowed in and the percentage of molecules in high FRET as a function of time was measured. **e,** This control was performed to verify that while performing loop-seq, molecules do not significantly unloop during the 20 minutes of digestion with RecBCD (see methods). If they do unloop, they would be immediately digested, which would affect the measurement of cyclizability. Sequence 5 (Supplementary Note 1) was used in this experiment. We find that digestion buffer (without the RecBCD enzyme) is in itself capable of looping molecules, and that too at a slightly faster rate than in the presence of looping buffer (which has 1M NaCl, see methods). This is owing to the presence of $Mg^{2+}$ ions in the digestion buffer, which we know to also effect looping by allowing stable hybridization of the ends[1]. Thus, molecules looped in the presence of 1M NaCl in 1 minute are expected to stay looped during the subsequent 20 minutes of digestion with RecBCD. **f,** This control was performed to understand the effect of the orientation of the central 50 bp sequence on the measured value of cyclizability. Two libraries were considered: the Random Library (where the sequence of DNA in the 50 bp central variable region are randomly selected – see Supplementary Note 5) and the Reverse Complement of the Random Library, where for every sequence, the left and right adapters are identical to that in the Random Library, whereas the central 50 bp is flipped (Supplementary Note 6). The cyclizabilities of sequences in the Random Library and the Reverse Complement of the Random Library are correlated. **g,** This plot serves to confirm the expectations that RecBCD does not digest looped molecules, and that over sufficient time, most molecules, even rigid ones, will loop. During timecourse loop-seq (Supplementary Note 3), the original sample was split into 8 identical fractions. Looping for various amounts of time and subsequent digestion was carried out for 7 of the 8 fractions, while one fraction was not subject to any digestion. All fractions were then PCR amplified (16 cycles) under identical conditions (see methods). Plotted are the concentrations of DNA obtained after PCR vs the corresponding times the samples were subject to the looping condition. These data points were fit to an exponential curve (solid line). The [DNA] obtained when no digestion was performed is represented as the dashed horizontal line. The fact that the fitted exponential approaches the dashed line indicates that for very long looping times, almost all molecules, even very rigid ones, have had sufficient time to loop and hence are protected from digestion. Thus, in this case, the concentration of DNA obtained after PCR of all surviving molecules approaches that of the fraction where no digestion was performed at all. Whether RecBCD would digest molecules sealed via non-canonical parallel basepairing or other geometries (Extended Data Fig. 1) is not known. However, this control suggests that either it does not, or such unconventional basepairings are rare. **h,** These plots serve to further demonstrate that if the library is permitted to loop for a very long time, most molecules, even very rigid ones, will loop, and that looped molecules are protected from subsequent digestion with RecBCD. In this case, the relative populations of various sequences measured after digestion should be similar to the case where no digestion was performed at all. We find this to indeed be the case: there is good correlation between the relative population of a sequence in sample 1 of timecourse loop-seq (no digestion) and sample 8 (2 hours of looping followed by digestion). However, correlation between the relative population of a sequence in sample 1 and that of a sequence in sample 2 (1 minute of looping prior to digestion) is much weaker. This is because in sample 2, only those molecules whose sequences render them bendable enough to loop under 1 minute are protected from subsequent RecBCD digestion. **i,** Technical replicates of the looping kinetic curve of sequence number 3 (Supplementary Note 1) measured using smFRET.

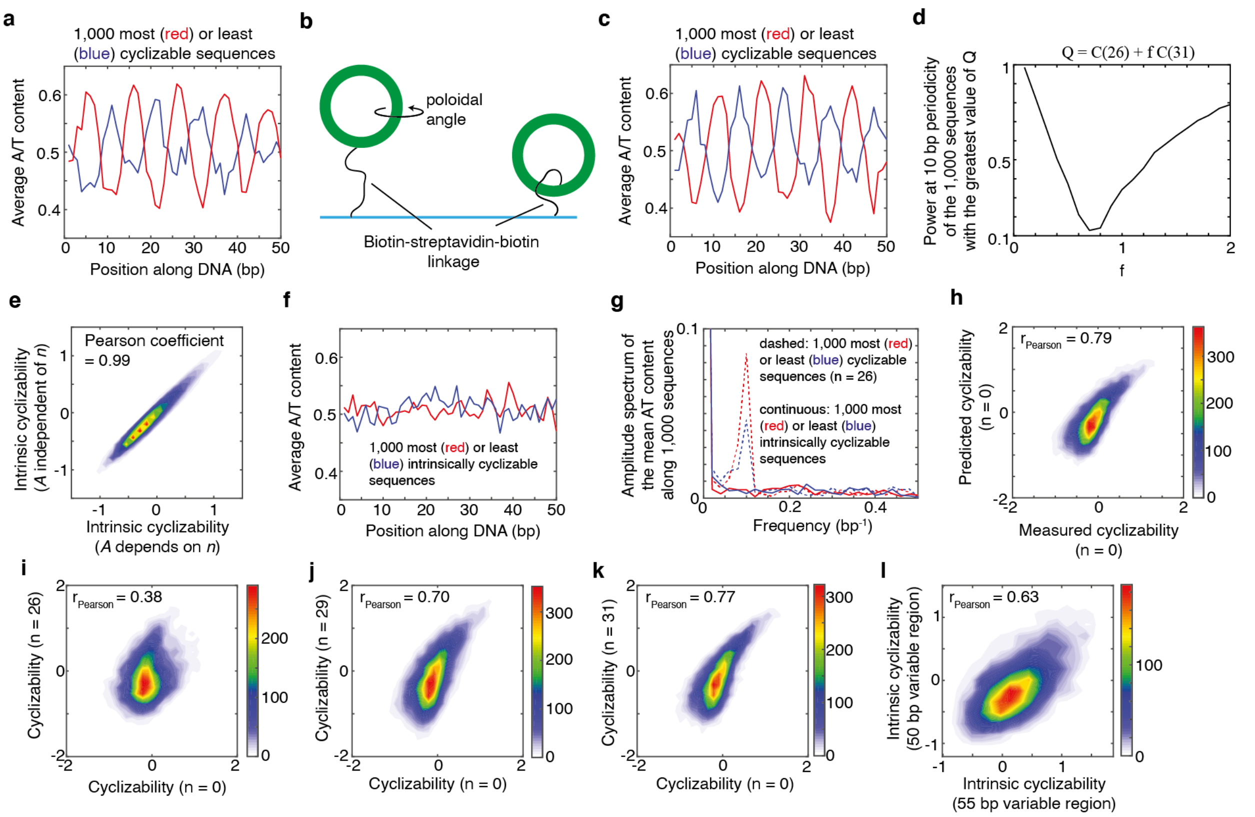


**Extended Data Fig. 4: Dependence of cyclizability on tether geometry and rotational phasing. a,** Loop-seq was performed on the Random Library (Supplementary Note 5). Plotted is the mean A/T content as a function of position along the central variable 50 bp region, where the mean is calculated by averaging over the 1,000 most cyclizable (red) or least cyclizable (blue) sequences. The value of n (n = 26) is the distance in nucleotides of the biotin tether from the end of each molecule in the library (Fig. 1d). **b,** In an untethered geometry, sequence features such as the phase of the oscillations in A/T content may result in the looped configuration having a preference for a certain poloidal angle (rotation along the long axis of DNA). Preference for a certain poloidal angle translates to a preference for a certain orientation of the biotin-streptavidin tether. Shown above are two extreme cases – in one case, the poloidal angle preference of the sequence results in a preferential orientation of the tether on the outside, while in the other case, the tether points to the inside at the point of contact with the DNA. As the biotin-streptavidin-complex is quite large, the outside orientation may be more favored for looping owing to steric considerations. The outside orientation can be converted to the inside orientation by moving the biotin tether point to a base that is half the DNA helical repeat away. This may explain why the phase of the oscillation of A/T content among the most or least cyclizable sequences shifts by half the helical repeat of DNA when the tether point is also shifted by half the helical repeat of DNA (about 5 bases) (compare panels a and c). **c,** The random library was re-prepared, placing the biotin 31 nt away from the ends (n = 31) and loop-seq was performed. Plotted are the same quantities as in panel a, except the 1,000 most and least cyclizable sequences of the library were identified based on the newly obtained cyclizability values under the n = 31 condition. **d – e,** See context in which these panels are referred to in Supplementary Note 7. **f,** Mean A/T content as a function of position along the variable region of the Random Library, where the averaging is done over the 1,000 sequences that have the highest (red) or lowest (blue) values of intrinsic cyclizability. The scale of the axes is the same as in panels a and c. **g,** Amplitude spectra obtained from the fast Fourier transforms of the plots in panel f (solid lines) and panel a (dashed lines). **h,** 2D histogram of the scatter plot of measured cyclizability of sequences in the Random Library prepared with the biotin at the very end of the molecule (n = 0 condition) vs its predicted value based on the oscillatory model (equation 1 in Supplementary Note 7). **i – k,** 2D histogram of scatter plot of measured cyclizabilities of sequences in the Random library prepared at n = 0 vs prepared at n = 26, 29, 31 nt. **l,** The use of long 10 nt overhangs has been shown to eliminate the need for ligase and to reduce the dependence of looping on rotational phasing between the ends[1]. Shown here is a 2D histogram of the scatter plot of intrinsic cyclizability of a sequence in the random library (which had 50 bp of DNA along the central variable region) vs the corresponding sequence in library L (Supplementary Note 8) where 5 bases were added to the variable region. A correlation coefficient only slightly poorer than the correlation between cyclizability values of the Random Library and the Reverse Complement of the Random Library (Extended Data Fig. 3f) suggests that rotational phasing of the ends does not significantly influence intrinsic cyclizability. See Supplementary Note 8.

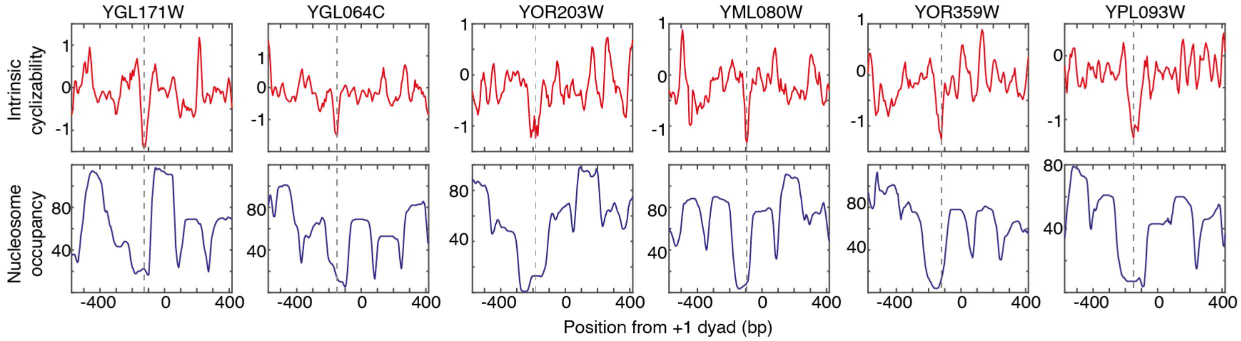


**Extended Data Fig. 5: Intrinsic cyclizability and nucleosome occupancy vs position from the dyads of the +1 nucleosomes of various individual genes in *S. cerevisiae*.** Plots are as shown for the two individual genes in Fig. 2c. The dashed line marks the ordinate value where intrinsic cyclizability is lowest.

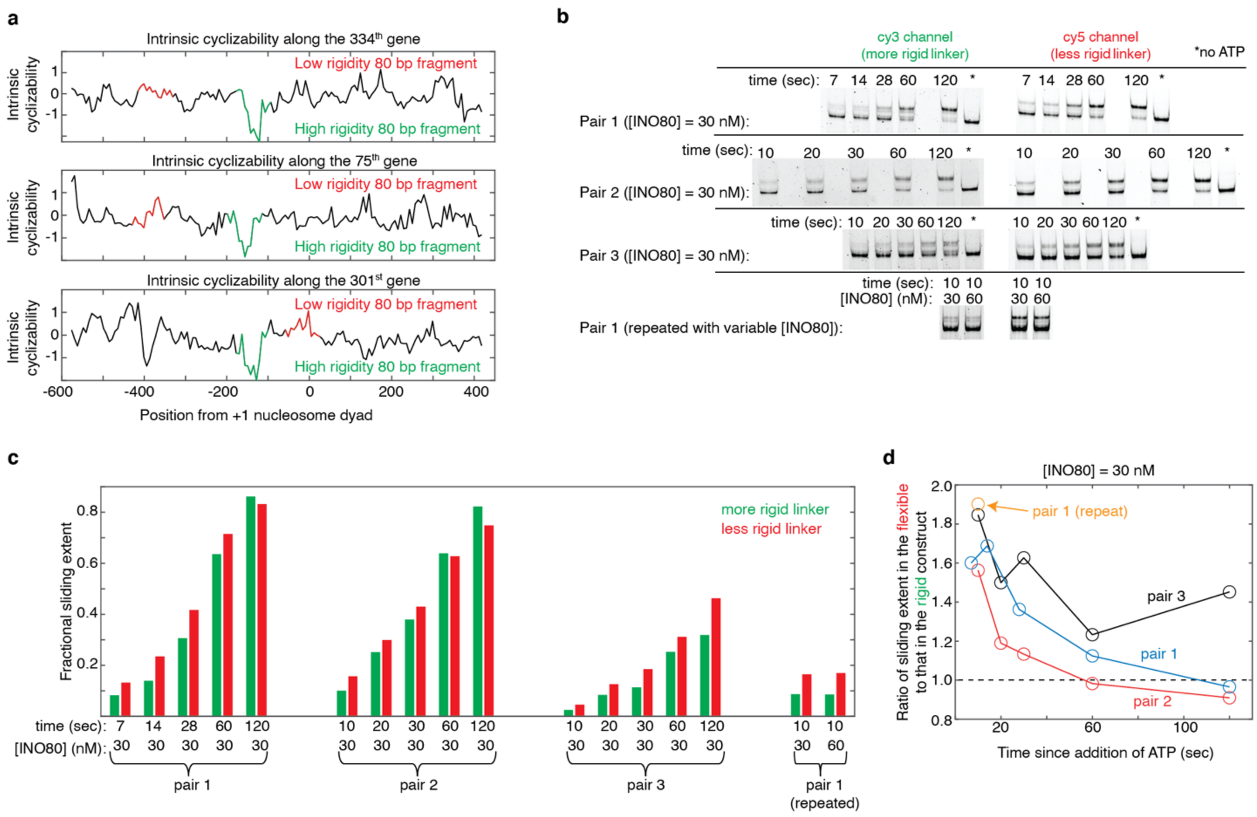


**Extended Data Fig. 6: DNA pair selection and timecourse of INO80 remodeling. a,** Intrinsic cyclizability as a function of position from the +1 nucleosomal dyad, along the 334th, 75th, and 301st genes in the list of 576 genes along which intrinsic cyclizability was measured (Supplementary Note 9). The 80 bp linker regions of both constructs (with rigid and flexible linkers extending from the 601 sequence) in pairs 1, 2, and 3 along which INO80 sliding extent was measured, were selected from the 334th, 75th, and 301st genes respectively. Genes are oriented in the upstream to downstream direction. Red and green denote the selected 80 bp less rigid and more rigid linker regions respectively. See Supplementary Note 11. **b,** The remodeling reactions shown in Fig. 2f were all performed for 1 minute of remodeling, under various enzyme concentrations. Here we show, instead, remodeling reaction timecourses at saturating [INO80] for all three pairs. Conditions are identical to that used in Fig. 2f, except that saturating (30 nM) INO80 is used and remodeling is permitted to progress for various amounts of time after addition of ATP (see methods). In all cases, the two constructs in a pair are present simultaneously, and distinguished by imaging the gel once for Cy3 and once for Cy5 fluorescence. Thus, although the sliding extent can be very sensitive to sliding time (especially for short sliding times), robust comparisons of sliding extents can be made between the two constructs in a pair. Quantification is done as explained in supplementary note 11. Sliding on the constructs in pair 1 was repeated in a separate experiment in presence of 30 nM enzyme, and also side by side in presence of 60 nM enzyme. Near identical extents of sliding indicate saturation has been reached at 30 nM INO80. **c,** Ratio of the fractional sliding extent in the construct formed on the more flexible linker to that formed on the more rigid linker, at various timepoints since addition of ATP, and in presence of 30 nM INO80. The dashed line indicates a ratio of 1. The ratio is computed from the data in panel b. The extent of sliding under saturating enzyme conditions is consistently higher for the construct involving more flexible linker (except, as expected, when the sliding extent approaches 100%). Solid lines connect observations that were made from the same initial reaction volume by sampling its fractions at various timepoints.

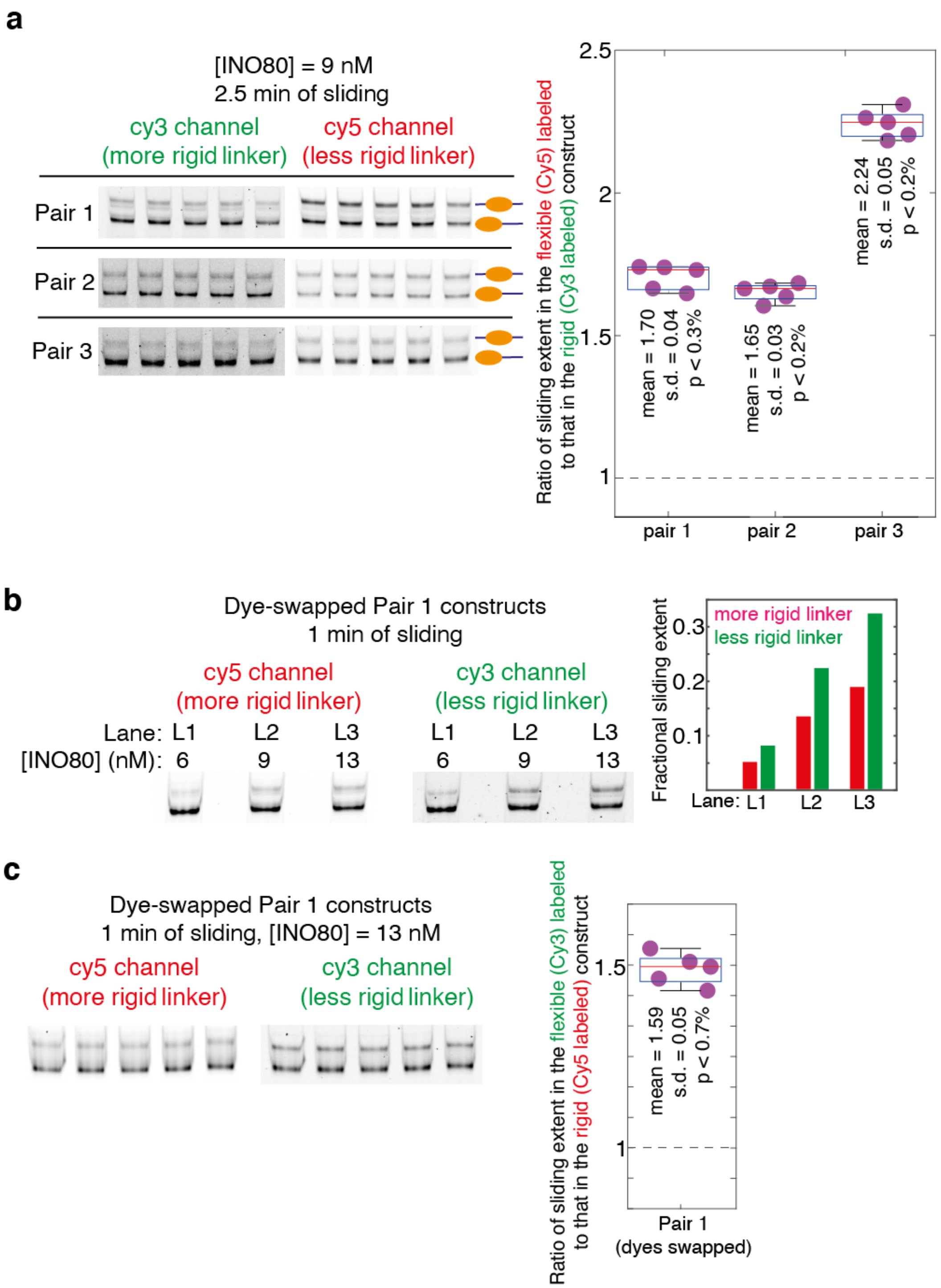


**Extended Data Fig. 7: Control experiments pertaining to the INO80-mediated sliding of nucleosomes as reported in Fig. 2f. a,** In order to assess our confidence in the result that INO80 mediated sliding is greater in the construct with the less rigid linker, we performed nucleosome sliding experiments similar to those reported in Fig. 2f five times for each pair in the presence of 9 nM INO80 and for 2.5 mins of sliding.

The products of sliding were analyzed on a 6% TBE gel as done in Fig. 2f. Each gel was imaged separately for Cy3 and for Cy5 fluorescence and quantified to calculate the fold-difference in sliding extent between the flexible and the rigid construct in each pair. The measurements of fold-differences for each pair are displayed in the box plots, along with the actual data points. Also indicated are the mean, standard deviation (s.d.), and the upper limit of the p value (defined here as the probability of obtaining a fold-difference of 1 if the distribution of fold differences has the same mean and s.d. as that of these 5 measurements) as obtained by the application of Chebyshev's inequality. Dashed line represents a fold-difference of 1. **b,** In the experiment described in Fig. 2f, the more rigid construct in all pairs was labeled with Cy3, while the less rigid construct was labeled with Cy5. This control verifies that the result that sliding extent in greater in the less rigid construct is not influenced by different dye properties. We swapped the dyes between the two constructs in pair 1. We then performed nucleosome sliding experiments on this modified pair 1 constructs for the three INO80 concentrations that yielded detectible sliding in Fig. 2f (6, 9, 13 nM), and for 1 minute of sliding as done in Fig. 2f. The products of sliding were analyzed on a 6% TBE gel, and the sliding extents quantified as done in Fig. 2f. We indeed find that even when the dyes are swapped, sliding extent is greater for nucleosomes formed on the less rigid construct. **c,** To obtain better statistics of sliding along the dye-swapped pair 1 constructs, we repeated one of the conditions in panel b (13 nM INO80, 1 min of sliding) five times. The measured fold-difference values are displayed in the box plot. Also indicated are the mean, standard deviation (s.d.), and the upper limit of the p value (defined as in panel a) as obtained by the application of Chebyshev's inequality. Dashed line represents a fold-difference of 1.

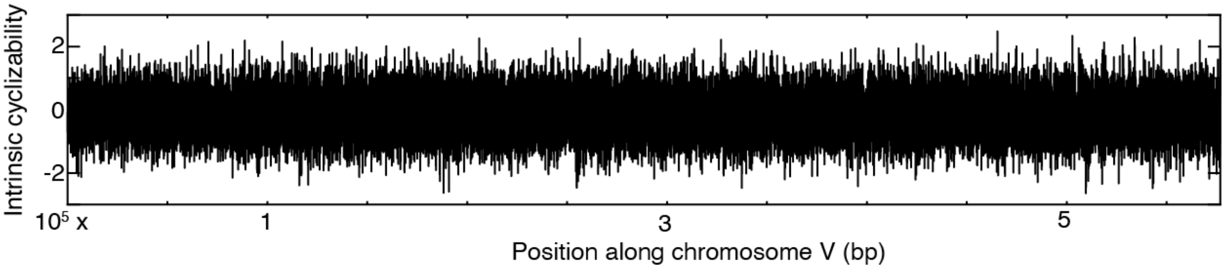


**Extended Data Fig. 8: Intrinsic cyclizability along *S. cerevisiae* chromosome V at 7 bp resolution.**

Data was obtained by performing loop-seq on the ChrV Library (Supplementary note 12).

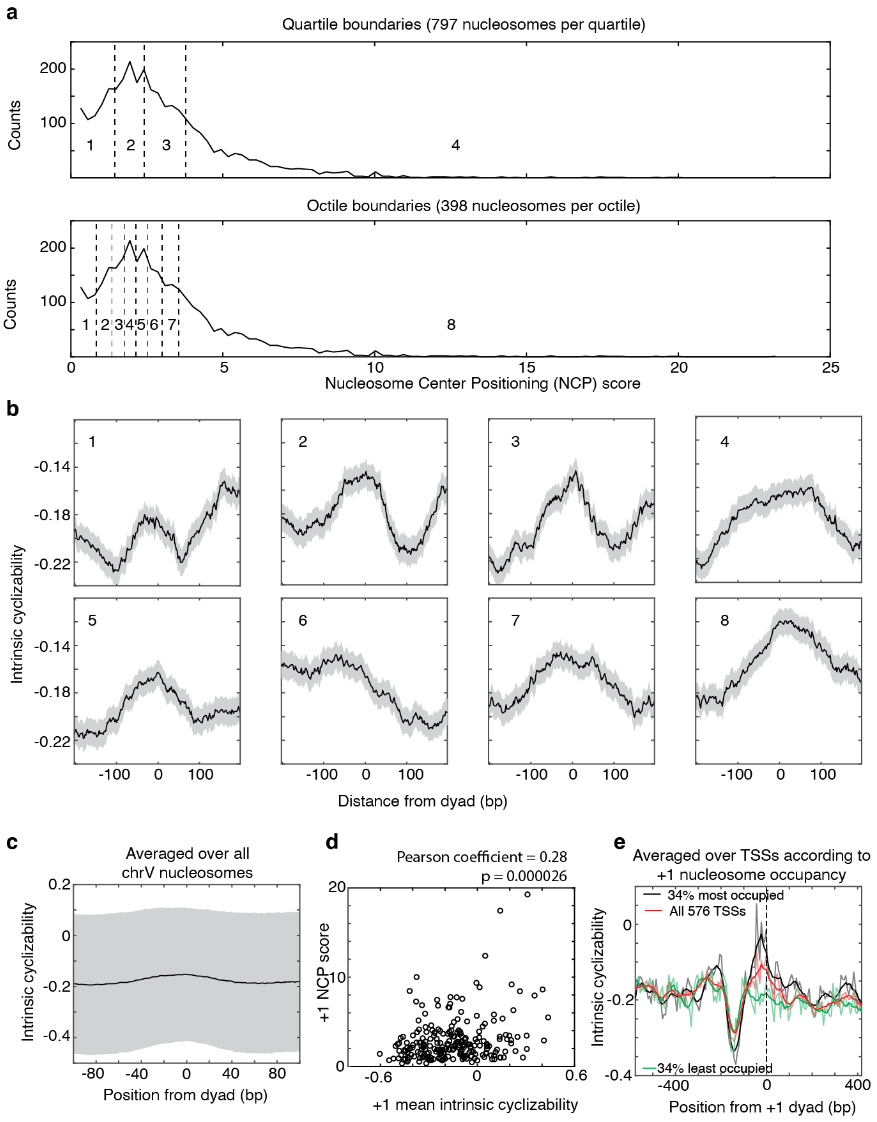


**Extended Data Fig. 9: Intrinsic cyclizability along nucleosomes. a,** Distribution of Nucleosome Center Positioning (NCP) scores[5] of all *S. cerevisiae* chromosome V nucleosomes. Quartile and octile boundaries of the distribution are shown in dashed lines and numbered (1 through 4 for quartiles and 1 through 8 for octiles). **b,** Mean intrinsic cyclizability of DNA as a function of position from the dyads of nucleosomes along yeast chromosome V, averaged over nucleosomes in each octile indicated in panel a. Error extents (shaded background) are s.e.m. **c,** Mean intrinsic cyclizability as a function of position, averaged over all *S. cerevisiae* chromosome V nucleosomes (solid line). Height of the shaded region is the standard deviation of measurements. **d,** Scatter plot of the NCP scores of the 227 +1 nucleosomes of the 227 genes identified along chromosome V vs the mean intrinsic cyclizabilities of DNA along the 147 bp that span these nucleosomes. Intrinsic cyclizability values were obtained by performing loop-seq on the ChrV Library. **e,** Plot of intrinsic cyclizability as a function of position along all the 576 genes in the Tiling Library (red), and among 34% of these genes that had the highest (black) or lowest (green) NCP score value of the gene's +1 nucleosome. Plots were obtained in a manner identical to that in Fig. 2b. Intrinsic cyclizability on either side of the dyad of +1 nucleosomes of genes that have high +1 nucleosome NCP score (black) is asymmetric, being higher on the TSS proximal (i.e. "left") side of the dyad.

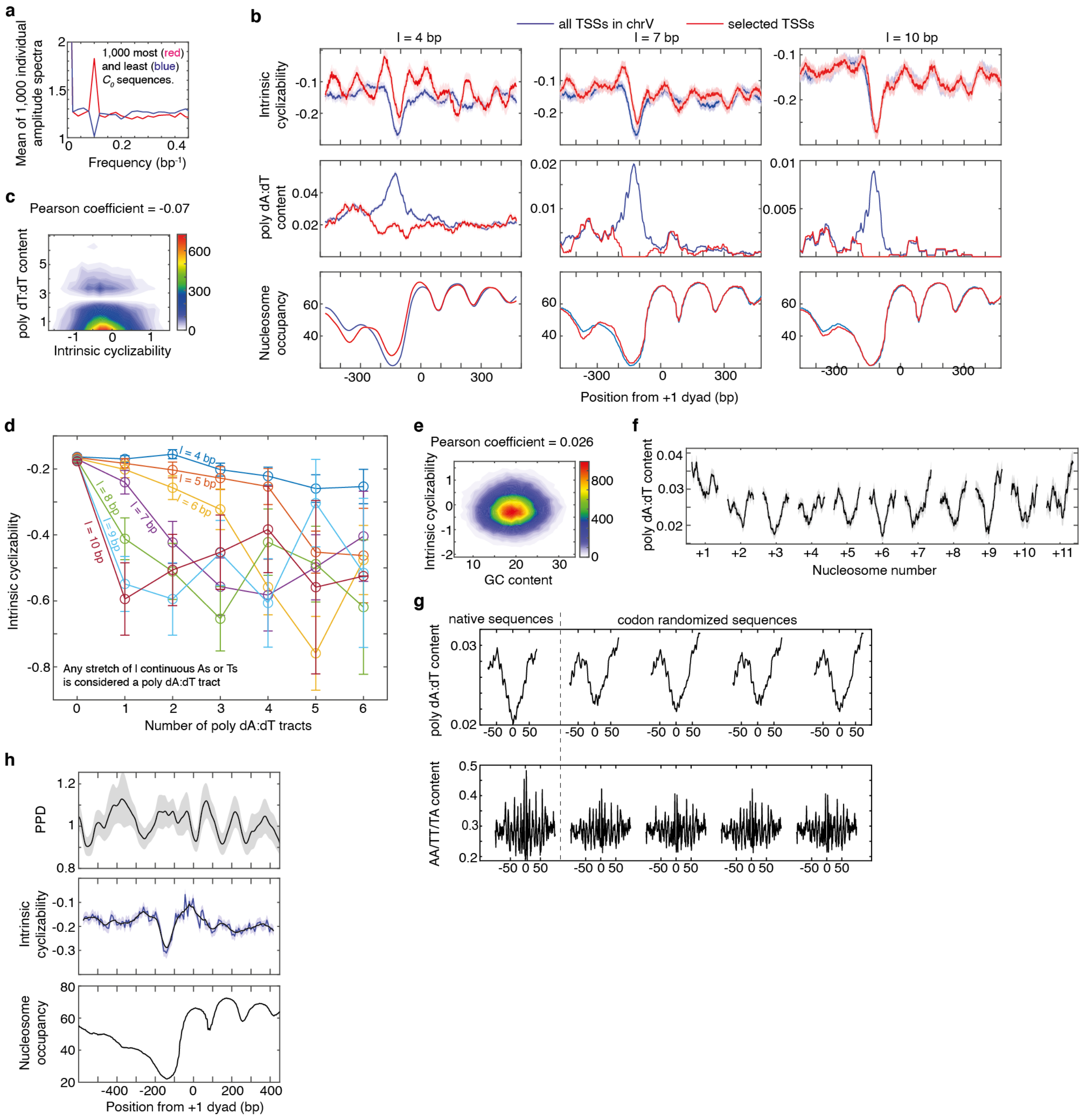


**Extended Data Fig. 10: Loop-seq measurements compared to expectations based on earlier measurements and models (see Supplementary Note 16). a,** Two sets of 1,000 plots each of A/T content as a function of position along the central 50 bp variable region of 1,000 sequences in the Random Library with the highest and lowest values of intrinsic cyclizability were generated. Fast Fourier transforms of these two sets of 1,000 plots were taken individually and used to calculate a total of 2,000 amplitude spectra. Plotted is the mean of the 1,000 amplitude spectra for the 1,000 sequences that have the highest (red) or lowest (blue) intrinsic cyclizability values. The plot indicates that sequences that have very high or low intrinsic cyclizabilities also tend to be characterized by enhanced or suppressed periodic modulations in AT content respectively at the DNA helical repeat. **b,** We consider a poly dA:dT stretch which has at least $l$ consecutive A or T nucleotides to be a poly dA:dT tract. For various values of $l$, plotted are intrinsic cyclizability (top panel), poly dA:dT tract content (middle panel), and nucleosome occupancy (bottom panel) vs position from the dyad of the +1 nucleosome, averaged over all 227 identified genes in *S. cerevisiae* chromosome V (blue) or over a selected subset of genes that show no peak in poly dA:dT content at the NDR (red; 30%, 62%, and 86% of genes for $l$ = 4 bp, 7 bp, 10 bp respectively). See supplementary note 15 for plotting details, including how poly dA:dT content is defined. **c,** 2D histogram of the scatter plot between the number of poly dA:dT tracts in the 50 bp variable region and intrinsic cyclizability of sequences in the ChrV library. Any stretch of $l$ or more consecutive As or Ts (here $l$ = 4) is considered a poly dA:dT tract. Thus a sequence with one stretch of 5 As, and no other As or Ts, in the 50 bp variable region has 2 poly dA:dT tracts if $l$ is considered to be 4. The scatter plot indicates that the overall correlation between intrinsic cyclizability and poly dA:dT content is very poor. Only non-overlapping sequences in the chrV library were considered. **d,** Binned histogram of the data in panel c (which represents the $l$ = 4 bp case), as well as for more restrictive definitions of poly dA:dT stretches ($l$ = 5 to 10 bp). The y-axis values are the mean intrinsic cyclizabilities of those sequences in the ChrV library which contain the number of poly dA:dT tracts in the central 50 bp variable region as specified along the x-axis. Error bars are s.e.m. **e,** 2D histogram of the scatter plot of mean GC content along the central 50 bp variable region of sequences in the ChrV library vs their intrinsic cyclizabilities. **f,** A plot of mean poly dA:dT content ($l$ = 4) as a function of position around the dyads of gene body nucleosomes along chromosome V in *S. cerevisiae*. The points along the horizontal axis where the nucleosome categories (+1, +2, etc) are marked represent the dyads of the nucleosomes. Light shaded region represents s.e.m. Poly dA:dT content was calculated as described in Supplementary Note 15. **g,** Mean poly dA:dT content ($l$ = 4) and AA/TT/TA vs position along the native and codon-randomized nucleosomal DNA sequences of the same 500 +7 nucleosomes along which intrinsic cyclizability profiles are reported in figure 3g. **h,** Predicted Plectoneme Density (PPD, see Supplementary Note 16) (top panel), intrinsic cyclizability (middle panel) and nucleosome occupancy (bottom panel) vs position from the +1 nucleosomal dyad, averaged over all 576 genes in the Tiling Library (supplementary note 9). PPD along each gene was first smoothened using a rolling window of 51 bp. The smoothened PPDs were then averaged at each position across all 576 genes, normalized by the mean, and plotted as the solid line in the top panel. The shaded background represents s.e.m. Intrinsic cyclizability and nucleosome occupancy were plotted exactly as in figure 2b. For the plot of intrinsic cyclizability, the solid blue and black lines represent data without and with a 7-fragment smoothening respectively (as in Fig. 2b). The shaded blue background represents s.e.m of intrinsic cyclizability values in the unsmoothed data.
