## Supplementary notes 1 - 18 for "Measuring DNA mechanics on the genome scale"

**Supplementary Note 1: generation of loopable molecules**

Our basic strategy to generate loopable molecules, either from a single template to obtain a looping kinetic curve via smFRET (Fig. 1a), or from a library to measure cyclizabilities via loop-seq (Fig. 1e) is described below.

**Brief overview:**

The initial templates are 100 nt single-stranded fragments. They are PCR amplified by primers that are complementary to the 25 bases at each end. In cases where a large number of templates in a library are amplified together, emulsion PCR (ePCR) is used instead of regular PCR to prevent incorrect annealing via the identical 25 bp at the ends. The primers also add 10 extra nts to either end of every molecule, and a biotin molecule close to one end. The amplified 120 bp DNA molecules are immobilized on neutravidin coated quartz slides (smFRET experiments) or streptavidin-coated beads (loop-seq). Recognition sites for the site-specific nicking enzyme Nt.BspQ1 are already built into the sequences of all template molecules near the ends. The molecules are nicked 10 nt from either end and the small 10 nt fragments still hybridized are washed away with buffer at 50 °C (Fig. 1b, see methods). This generates immobilized DNA molecules with a central 100 bp duplex region, flanked by complementary 10 nt single-stranded overhangs.

**Detailed examples:**

The following are the 10 templates that we used to generate the 10 DNA molecules whose individual looping kinetics were measured using smFRET (Fig. 1c). The 25 bp red regions that flank the central 50 bp black region are identical among the seven fragments. The black region is variable.

Sequence 1:
TTTCTTCACTTATCTCCCACCGTCCGGAATGGTTACCGAAAACGGGCTTCTTCTCATCCATATCCATAAT
TTCATGGCAGAAGACAAGGGAACGAAATAG

Sequence 2:
TTTCTTCACTTATCTCCCACCGTCCTTTGTCAGGGTGTTCTTCGATTTTCCTTAGGTGCTTTTAGGACTT
GTTTAGGCAGAAGACAAGGGAACGAAATAG

Sequence 3:
TTTCTTCACTTATCTCCCACCGTCCTGTTACGCATACACACAAGTACAGATTTGTATATAGTATTTCTTC
TTCGTGGCAGAAGACAAGGGAACGAAATAG

Sequence 4:
TTTCTTCACTTATCTCCCACCGTCCTGCTAATCCATATCATATAATCTAAAAACAAATCTAAAGGATCAT
CCATAGGCAGAAGACAAGGGAACGAAATAG

Sequence 5:
TTTCTTCACTTATCTCCCACCGTCCAGGGCTTTTCTGTATTTTCTCATAGACATTATTTATCAGTAATTG
CAGCTGGCAGAAGACAAGGGAACGAAATAG

Sequence 6:
TTTCTTCACTTATCTCCCACCGTCCCAAAAGTGACATCCACAGCAAGCTGGACAGGTAAATTGCCTCATA
CAATCGGCAGAAGACAAGGGAACGAAATAG

Sequence 7:
TTTCTTCACTTATCTCCCACCGTCCCTTCAACGCACCTTAAATCTATTCTTTTAAATTTTCCAGATTCTA
TTAACGGCAGAAGACAAGGGAACGAAATAG

Sequence 8:
TTTCTTCACTTATCTCCCACCGTCCTTTATCTCAAATTTGCGCTTTTTAGTCCAATCTCTCACAGTGACT
ATAGTGGCAGAAGACAAGGGAACGAAATAG

Sequence 9:
TTTCTTCACTTATCTCCCACCGTCCCGACTATTTTTTAATTGATAGTTCTGAAGGCTCTTCTACTGATGA
CGAACGGCAGAAGACAAGGGAACGAAATAG

Sequence 10:
TTTCTTCACTTATCTCCCACCGTCCTGTCTACTTGAAATTCTAATTCATATTTTTTTTGTTTGGATAGAA
TATCAGGCAGAAGACAAGGGAACGAAATAG

The above templates were PCR amplified using the primers P1 and P2 listed below 5' to 3':

P1:/5Cy3/CAGAATCCGTCGAAGAGCCTTATCTCCCACCGTCC

P2:/5Cy5/ACGGATTCTGCGAAGAGCTTCCCTTG/iBiodT/CTTCTGCC

Here, /5Cy3/ and /5Cy5/ represent 5' attachment of the fluorophores Cy3 and Cy5, respectively. The fluorophores are included only for smFRET experiments and are absent for molecules that are used in the loop-seq assay. /iBiodT/ represents a biotin attached to a modified thymine base. Blue indicates the region that is complementary to the ends of the template DNA sequences. These primers also serve as the primers used to amplify all subsequent libraries on which loop-seq was performed.

As an example, when Sequence 1 is PCR amplified, the following 120 bp product is obtained:

```
5' - Cy3 CAGAATCCGTCGAAGAGCCTTATCTCCCACCGTCCGGAATGGTTACCGAAAACGGGCTTC...
     3' - GTCTTAGGCAGCTTCTCGGAATAGAGGGTGGCAGGCCTTACCAATGGCTTTTGCCCGAAG...

 ...TTCTCATCCATATCCATAATTTCATGGCAGAAGACAAGGGAAGCTCTTCGCAGAATCCGT - 3'
 ...AAGAGTAGGTATAGGTATTAAAGTACCGTCTTCTGTTCCCTTCGAGAAGCGTCTTAGGCA Cy5 - 5'
```

Here, the central 50 bp variable region is in black, the adapters on either end identical among all molecules are in blue, the recognition sequence for the nicking enzyme Nt.BspQ1 are in red, and the Thymine on which the biotin moiety is attached is in brown.

Following immobilization through biotin, nicking by Nt.BspQ1, and washing with buffer at 50 $^{o}$C (Fig. 1b), the molecule is converted to an immobilized loopable molecule with a central 100 bp duplex region flanked by 10 nt complementary single-stranded overhangs (Fig. 1d):

```
5' - Cy3 CAGAATCCGTCGAAGAGCCTTATCTCCCACCGTCCGGAATGGTTACCGAAAACGGGCTTC...
               3' - GCTTCTCGGAATAGAGGGTGGCAGGCCTTACCAATGGCTTTTGCCCGAAG...

...TTCTCATCCATATCCATAATTTCATGGCAGAAGACAAGGGAAGCTCTTCG - 3'
...AAGAGTAGGTATAGGTATTAAAGTACCGTCTTCTGTTCCCTTCGAGAAGCGTCTTAGGCA Cy5 - 5'
```

**Supplementary Note 2: definition of cyclizability**

Cyclizability is defined is the natural logarithm of the ratio of the relative population of a sequence in the selected pool to that in an identically treated control pool where no digestion was performed (Fig. 1e). Below we relate cyclizability to other quantities such as the probability of looping and the energy barrier for the looping transition.

A typical number of total molecules in the sample pool is ~ $6 \times 10^{10}$. As this is a very large number compared to the number of different sequences (~ 10,000 – 90,000), we assume after splitting the sample identically by volume into two fractions (Fig. 1e), that for every sequence, an equal number of copies exists in each of the two fractions.

If $n_i$ is the number of copies of the $i^{\text{th}}$ sequence in either pool prior to looping, and $p_i$ is the probability that the $i^{\text{th}}$ sequence will loop in under 1 minute, then cyclizability of the $i^{\text{th}}$ sequene, $C^i$, as defined, is given by:

$$C^i = ln\left( \frac{p_i n_i}{\sum_i p_i n_i} \Big/ \frac{n_i}{\sum_i n_i} \right) = \ln p_i + \ln \frac{\sum_i n_i}{\sum_i p_i n_i} = \ln p_i + C \text{ ------------ (1)}$$

where $C$ is independent of $i$ (it is summed over all $i$). $C$ is thus an additive constant that is added to the ln $p_i$ term of every sequence. However, the value of this constant is specific to the library in question, which is why cyclizability values of sequences belonging to different libraries can only be compared up to an additive constant.

The use of natural logarithm permits relating cyclizability to looping energy as follows:

An Arrhenius model for the looping rate $k_l$ is given by:

$$k_l = Ae^{-\frac{E_a}{k_B T}} \text{ ---------------- (2)}$$

where $A$ is a frequency factor, $E_a$ is the barrier height for the looping transition, and $k_BT$ is the thermal energy. As looping curves typically fit to single exponentials we assume:

$$p_i = 1 - e^{-k_l T_l} \text{ -------------- (3)}$$

where $T_l$ is the time for which looping is permitted ($T_l$ = 1 min for most experiments).

Using timecourse loop-seq (Supplementary Note 3) performed on a library comprising 19,907 DNA fragments ('Cerevisiae Nucleosomal Library', Supplementary Note 4), we find that looping times (i.e., inverse of looping rates) have a typical median of 12.3 mins, and an inter-quartile range of 14 mins. Only 1.4% of sequences have a looping time less than 1 minute. Thus $k_lT_l$ for $T_l$ = 1 min is typically less than 1. Thus under these conditions, an approximate expression for $p_i$ is given by:

$$p_i = 1 - e^{-k_l T_l} \approx 1 - (1 + k_l T_l) = k_l T_l \text{ ------------ (4)}$$

Thus using (1) and (2):

$$C^i \approx -\frac{E_a}{k_B T} + D \text{ ------------- (5)}$$

when $k_l T_l$ <<1, which we show is satisfied by most sequences in a large library of genomic sequences. $D$ is a constant term which is specific to the library but not to individual sequences within the library. Nevertheless, we do not explicitly relate cyclizability of a sequence to its barrier energy for the looping transition because cyclizability is sufficient to make a relative comparison of the looping propensity among sequences in a library.

**Supplementary Note 3: timecourse loop-seq**

In timecourse loop-seq, a single large sample of DNA bound to beads was processed as described for regular loop-seq (methods). Just prior to looping in high salt, the sample was split into 8 equal volumes, each ~95 µl. The beads in samples 1 - 7 were allowed to loop for 1, 5, 10, 15, 20, 40, 120 minutes respectively, followed by digestion of unlooped molecules. Beads in sample 8 were never digested. Then, as described in the protocol for loop-seq (see methods), beads in all 8 samples were resuspended in high salt looping buffer, and amplified via PCR for 16 cycles. PCR mixtures were purified using Ampure beads (Agencourt) and the concentration of DNA obtained for each of these samples was measured using Qubit. The samples were then sequenced, and the relative population of each sequence in each of the 8 samples was calculated.

For the $j^{\text{th}}$ sequence, $[t_i, y_i]$ where $y_i = c_i r_i^j$ for $i = 1,2, \ldots ,8$ represents a set of 8 data points, where $t_i$ is the time for which the $i^{\text{th}}$ sample was looped, $c_i$ is the total concentration of recovered DNA in the $i^{\text{th}}$ sample as measured via Qubit, and $r_i^j$ is the relative population of the $j^{\text{th}}$ sequence in the $i^{\text{th}}$ sample as measured via Illumina sequencing. This set of 8 data points was fit to the equation $y = A(1 - e^{-t/\tau})$. $\tau$, as obtained from the fit, is the looping time of the $j^{\text{th}}$ sequence.

See Extended Data Fig. 2 for a comparison between looping kinetic curves obtained using individual smFRET experiments and using timecourse loop-seq, and for a scatter plot of cyclizability (defined for 1 minute of looping only) vs. looping energy barrier (natural logarithm of the looping time, as obtained from timecourse loop-seq, supplementary note 2). Because this latter plot is essentially linear, we focused on only measuring cyclizability for subsequent loop-seq runs.

**Supplementary Note 4: construction of the Cerevisiae Nucleosomal Library**

This library comprised 19,907 different sequences. The dyad locations (in the SacCer2 version of the *S. cerevisiae* genome assembly) of the 10,000 nucleosomes among all reported nucleosomes with the highest Nucleosome Center Positioning (NCP) scores[1] were selected. The sequences of the 50 bp DNA fragments immediately to the left and right of the dyads of these nucleosomes were noted (SacCer2 assembly). The dyad itself was not included. 19,907 sequences from among these 20,000 sequences were selected to constitute the central 50 bp variable region of all sequences in this library. They were flanked by the standard 25 bp adapters and 10 nt overhangs described in Supplementary Note 1.

### Supplementary Note 5: construction of the Random Library

This library comprised 12,472 DNA sequences. Sequences in the central 50 bp variable regions were specified at random using a random number generator (MATLAB). Each nucleotide was equally likely to occur at any location in this region. This region was flanked by the standard 25 bp left and right adapters and 10 nt overhangs (Supplementary Note 1).

**Supplementary Note 6: construction of the Reverse Complement of the Random Library**

This library, like the random library, comprised 12,472 100 bp sequences. There is a one to one correspondence between the sequences in this library and sequences in the random library, as explained in the diagram below:

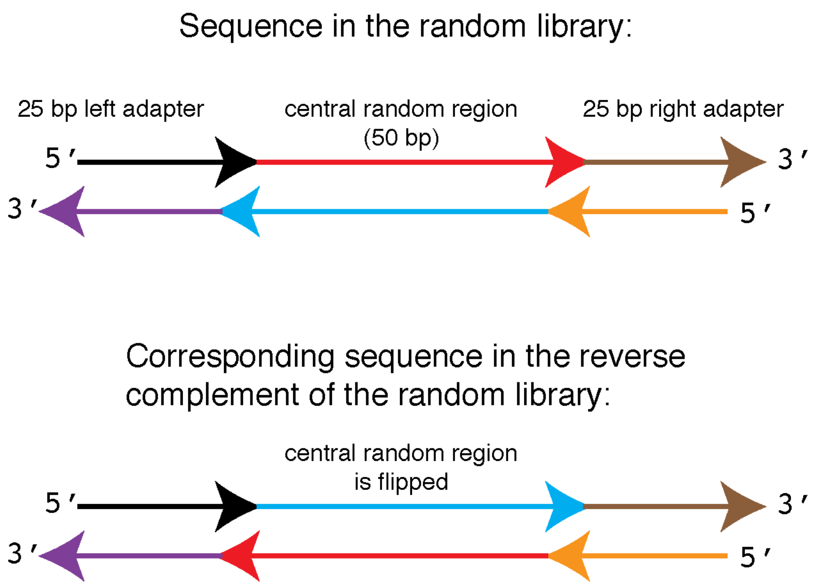

**Supplementary Note 7: the location of the biotin tether can influence DNA cyclizability**

**The biotin tether location influences cyclizability:**

While performing loop-seq on a library of 12,472 randomly generated DNA sequences ('Random Library', supplementary note 5), we noticed that the mean A/T content along the central 50 bp variable region of the 1,000 most cyclizable sequences oscillates with ~10 bp periodicity, which is about the helical repeat of DNA (Extended Data Fig. 4a). The 1,000 least cyclizable sequences also showed oscillatory mean A/T content, but out of phase compared to the most cyclizable sequences (Extended Data Fig. 4a). Although periodic oscillations in A/T content may be a feature of sequences that have extremely high or low cyclizabilities, there should not be a preferred phase as the sequences belong to the Random Library. This phasing effect reminded us of the observation that native nucleosomal DNA have AA/TT/TA dinucleotides enriched where the minor groove bends inwards[2,3], suggesting that it is due to a favored poloidal angle (rotation along the long axis of DNA, Extended Data Fig. 4b) for each sequence. We hypothesized that if the favored poloidal angle puts the biotin, i.e. the surface-tethering point, on the outer surface of the DNA circle, cyclizability is higher and vice versa (Extended Data Fig. 4b). Indeed, when we moved the biotin by 5 bp, about half a helical turn, i.e. from n = 26 to n = 31 with n as defined in Fig. 1d, the phase of A/T content oscillations at which looping is favored or disfavored also shifted by half the helical repeat (compare panels a and c in Extended Data Fig. 4).

**An oscillatory model to capture the influence of the tether location on cyclizability:**

In order to extract an intrinsic quantity that is independent of the tether location, we constructed a simple model sufficient to capture the tether position effects. We assumed an oscillatory dependence of cyclizability on the tether position with periodicity equal to the helical repeat:

$$C^i(n) = C_0^i + A^i \sin(kn + \varphi^i), \text{ where } k = \frac{2\pi}{10.4}\ bp^{-1} \text{ ------- (1)}$$

Here, $C^i(n)$ is the cyclizability of the $i^{th}$ sequence when the tether point is *n* nt away from the end (Fig. 1d). $C_0^i$, $A^i$ and $\varphi^i$ are the mean, amplitude, and phase terms for the $i^{th}$ sequence. We performed loop-seq three times with the biotin at three positions (n = 26, 29, 31) and measured $C^i(n)$ for n = 26, 29, 31. For each $i$, we solved the three equations obtained by plugging n = 26, 29, 31 into (1) and obtained $C_0^i$, $A^i$ and $\varphi^i$. $C_0^i$ is the mean term which we call the intrinsic cyclizability of the $i^{th}$ sequence.

**Refinements of the model:**

If $A^i$ has no dependence on $n$, the quantity $Q(f) = C^i(26) + fC^i(31)$ should have almost no dependence on $n$ when $f = 1$ because in this case, the two sinusoidal quantities being added are almost 180° apart in phase. For various values of $f$, we ranked the sequences in the Random Library in order of increasing $Q(f)$, and chose the 1,000 sequences with the highest values of $Q(f)$. We then plotted the mean A/T content as a function of position along the central 50 bp variable region of these 1,000 sequences, obtained the amplitude spectrum by taking the fast Fourier transform of the plot, and noted the power at 10 bp period. We found that $f = 1$ is not the point where this power is lowest, but rather it is at $f = 0.7$ (Extended Data Fig. 4d).

The simplest way to incorporate this observation into the model is to assign a dependence of $A^i$ on $n$ of the form:

$\frac{A^i(n=31)}{A^i(n=26)} = 0.7 for\ all\ i$ ------- (2)

By linear interpolation, we also enforce:

$\frac{A^i(n=29)}{A^i(n=26)} = 0.82\ for\ all\ i$ ------ (3)

We found that incorporating such a dependence of $A$ on $n$ only insignificantly changes the value of intrinsic cyclizability (Extended Data Fig. 4e). Nevertheless, all reported cyclizability values took into account a dependence of $A$ on $n$ as described in (2) and (3) above.

We also checked if the ratio of amplitudes as defined by (2) and (3) above depends on other sequences present in the library. We therefore combined the Cerevisiae Nucleosomal Library (which has 19,907 sequences, supplementary note 4) and the Random Library (which had 12,472 sequences, supplementary note 5) to create a library with 32,379 members. We measured the cyclizabilities of all sequences in this combined library at three biotin locations (n = 26, 29, 31). We then used the cyclizability values of the sequences that belonged to the random library part of this combined library to re-estimate the value of $\frac{A^i(n=31)}{A^i(n=26)}$ as described above, and still found it to be 0.7.

**$C_0$ (intrinsic cyclizability) is a quantity unaffected by tether location:**

Supplementary Data reports all three values for most sequences we analyzed in this study. The mean AT content along the central 50 bp variable region of sequences in the random library with very high or low values of $C_0$ (which we call intrinsic cyclizability) does not show any oscillations in AT content, indicating successful removal of the oscillatory modulations imposed by specific tether geometries, or for that matter, by any other factor such as the sequence of the flanking 25 bp left and right adapters[4] (Extended Data Fig. 4f-g).

**Testing the oscillatory model:**

We tested this oscillatory model by measuring cyclizability of sequences in the Random Library at a fourth biotin location (n = 0). The previously measured values of $C^i(n = 26)$, $C^i(n = 29)$ and $C^i(n = 31)$ were used to calculate $C_0^i$, $A^i$ and $\varphi^i$ using equation (1) for all sequences $i$ in the Random library. The values were then plugged back into equation (1) to predict $C^i(n = 0)$. A 2D histogram of the scatter plot between the predicted and measured cyclizabilities shows a good correlation (Extended Data Fig. 4h) that is, in fact, better than the correlation between cyclizability measurements of the Random Library and the Reverse Complement of the Random Library (Extended Data Fig. 3f).

Also, since the biotin tether location for the n = 0 case is almost an integer number of helical turns (10.4 bp/turn) away from the tether location in the n = 31 case, and an odd integer number of half helical turns away from the n = 26 case, we predicted $C^i(n = 0)$ to be best correlated with $C^i(n = 31)$ and least correlated with $C^i(n = 26)$. We indeed find this to be the case (Extended Data Fig. 4i–k), further confirming that the influence of the biotin tether location on DNA cyclizability can be captured by an oscillatory model with a period equal to the helical repeat of DNA.

**Supplementary Note 8: control for assessing the dependence of intrinsic cyclizability on rotational phasing**

Earlier works demonstrated that the use of long 10 nt overhangs allows stable hybridization of the ends during looping without the need to "seal" the looped configuration via ligase action[5]. This was further shown to significantly reduce the dependence of looping rate on the rotational phasing of the two ends[5]. We verified this to indeed be the case, by measuring the intrinsic cyclizabilities of a set of 6,144 sequences, and comparing them to the intrinsic cyclizabilities values obtained when the central 50 bp region was lengthened by about half the DNA helical turn to 55 bp. The first 6,144 sequences of the Random Library (Supplementary Note 5) were selected, and a new library was constructed where the central 50 bp variable region is preceded by a 5 bp stretch. These new sequences (which had 55 bp instead of 50 bp in the central variable region) were represented between sequence numbers 61,145 and 67,288 of library L (supplementary note 14).

First, all 1024 pentamers were listed cyclically as:

P1: AAAAA
P2: AAAAT
P3: AAAAG
P4: AAAAC
P5: AAATA
…
P1024: CCCCC

These 1024 pentamer sequences were cyclically added to the start of the variable regions of library members 1 – 6,144 of the Random Library. The resulting fragments were included in Library L (Supplementary Note 14). The following is a description of the central variable region (55 bp in length) of sequence numbers 61,145 – 67,288 of library L:

Sequence # 61,145 of library L is P1 + the central 50 bp region of library member 1 of the Random Library
Sequence # 61,146 of library L is P2 + the central 50 bp region of library member 2 of the Random Library
…
Sequence # 62,168 of library L is P1024 + the central 50 bp region of library member 1024 of the Random Library
Sequence # 62,169 of library L is P1 + the central 50 bp region of library member 1025 of the Random Library
…
Sequence # 67,288 of library L is P1024 + the central 50 bp region of library member 6144 of the Random Library.

See Extended Data Fig. 4l for the scatter plot of intrinsic cyclizability of a sequence in the random library (which had 50 bp of DNA along the central variable region) vs the corresponding sequence in library L where 5 bases were added to the variable region. This correlation coefficient is only a little poorer than the correlation between cyclizability values of the Random Library and the Reverse Complement of the Random Library (Extended Data Fig. 3f). One has to further, of course, consider that there is an actual difference in DNA sequence and length between the two quantities plotted here, which would be expected to cause a lower correlation even if rotational phasing has no effect.

**Supplementary Note 9: construction of the Tiling Library**

A set of 576 genes from *S. cerevisiae* were selected. To generate this list, all annotated genes that were described as ORFs and had both ends mapped with high confidence in a previous study[6] were considered. The first 297 genes in the list were chosen at random from among such genes, while the subsequent 279 genes were the most expressed genes among all genes we had identified. Expression level was defined as the mean number of RNA polymerases per base along the entire length of the transcripts, or along the first 500 bp if the length of the transcripts is greater than 500 bp. RNA polymerase counts were obtained from reported NETseq measurements[7].

The coordinates of the dyads of the +1 nucleosomes of these 576 genes were obtained from an earlier report[1] and converted to the SacCer3 assembly using liftover[8]. For every gene, a 2,001 bp region was selected, from the position (dyad – 1,000) to (dyad + 1,000) for genes on the positive strand and (dyad + 1,000) to (dyad – 1,000) for genes on the negative strand (according to the SacCer3 assembly). Here 'dyad' refers to the dyad of the +1 nucleosome of that gene. Therefore, irrespective of the strand, the 2,001 bp region was oriented in the upstream to downstream direction. From this 2,001 bp region, the following 143 50 bp fragments were selected:

Segment 1: position 400 to 449
Segment 2: position 407 to 456
…
Segment 143: position 1394 to 1443

This process was repeated for all 576 genes to obtain 82,368 fragments in the tiling library. The fragments were flanked by the standard 25 bp left and right adapters (Supplementary Note 1).

### Supplementary Note 10: plotting details for Fig. 2b

For every gene among the 576 genes, the Tiling Library (Supplementary Note 9) contained 143 50 bp DNA fragments, the center of each fragment being at a specific distance from the dyad of the +1 nucleosome of that gene (Fig. 2a). After measuring the intrinsic cyclizabilities of these fragments, we constructed a matrix of 576 rows and 143 columns. Each row represents a gene in the library, and its columns were populated with the cyclizability values of the 143 DNA fragments that tiled that particular gene. Plotted in grey is the column-wise average of this matrix. Plotted in red is the grey curve smoothened over a sliding window of 7 fragments. The 143 x-axis values represent the distances of the centers of the 143 fragments from the location of the +1 nucleosome dyad.

To obtain the plot of nucleosome occupancy as a function of position (bottom panel), we first obtained nucleosome occupancy at every genomic position along the genome of *S. cerevisiae* by re-analyzing published data, in a manner similar to what was done earlier[9]. Only the first replicate (GSM2561057) from GEO accession number GSE97290 was used. Reads were aligned to SacCer3 with bowtie2[10] using a maximum fragment length of 1,000 and "very-sensitive" preset options. From the aligned reads fragments of length in the range 44 – 58 bp were selected as nucleosomes. The centers of these fragments were considered as the nucleosomal dyads. For a give genomic position, the number of such identified nucleosomal centers found in reads that lie within 73 bp of it was counted. In other words, this count is the number of identified nucleosomes whose 147 bp extent covers this genomic location. This count was defined as the nucleosome occupancy. Having obtained nucleosome occupancy at every genomic location, a similar approach was used to plot the curve as was done for the top panel described above. The corresponding matrix was populated with nucleosome occupancy values instead of intrinsic cyclizability values and the resolution was higher because nucleosome occupancy values are available for every location (unlike intrinsic cyclizability which was measured every 7 bp).

### Supplementary Note 11: sliding of nucleosomes by INO80

#### Construct design:

Six different 227 bp DNA constructs were prepared. The first 147 bases of all six constructs were identical to the 601 DNA sequence, while the subsequent 80 bases were different among the six constructs. All six of the 80 bp regions were selected from various regions of the 576 genes along which we had measured intrinsic cyclizability (Tiling Library, supplementary note 9). The six constructs were grouped into three pairs such that between the two constructs in each pair, there was a large difference in the intrinsic cyclizability values near the middle of the 80 bp region (Fig. 2e).

The two 80 bp regions for the two constructs in pair 1 were both selected from the 334$^{th}$ gene.
The two 80 bp regions for the two constructs in pair 2 were both selected from the 75$^{th}$ gene.
The two 80 bp regions for the two constructs in pair 3 were both selected from the 301$^{st}$ gene.

Plotted in Extended Data Fig. 6a are intrinsic cyclizabilities along the 334$^{th}$, 75$^{th}$ and 301$^{st}$ genes (from the list of 576 genes). In each plot, marked in red and green are the regions that were chosen for the two 80 bp fragments, one with much higher rigidity (lower intrinsic cyclizability) in the middle of the 80 bp region (green) than the other (red).

Pair 1:

The sequence with low rigidity (high intrinsic cyclizability) in the 80 bp region is:

ACAGGATGTATATATCTGACACGTGCCTGGAGACTAGGGAGTAATCCCCTTGGCGGTTAAAACGCGGGGG
ACAGCGCGTACGTGCGTTTAAGCGGTGCTAGAGCTGTCTACGACCAATTGAGCGGCCTCGGCACCGGGAT
TCTCCAGAAAGTGCAAGTGAAGGAGTCAGAACTGCCCTCCTCTATACCGGCGCAGACTGGATTGACGTTT
AATATATGGTATAATAA

The sequence with high rigidity (low intrinsic cyclizability) in the 80 bp region is:

ACAGGATGTATATATCTGACACGTGCCTGGAGACTAGGGAGTAATCCCCTTGGCGGTTAAAACGCGGGGG
ACAGCGCGTACGTGCGTTTAAGCGGTGCTAGAGCTGTCTACGACCAATTGAGCGGCCTCGGCACCGGGAT
TCTCCAGCGTTTTTTACTTTTCTTTATTCCTCATCACTTTTTTTCAGAAAAATTTTTTTCAGTTTTTGAG
CATTCCATCGTTACATA

In both these cases, the 601 part of the sequence is denoted in black and the 80 bp segment is denoted in red. The 80 bp regions (red in the sequences above) for both these sequences were selected from among the 576 genes characterized earlier in Fig. 2b. See Extended Data Fig. 6a for plots of intrinsic cyclizability across these linker regions.

Pair 2:

The sequence with low rigidity (high intrinsic cyclizability) in the 80 bp region is:

ACAGGATGTATATATCTGACACGTGCCTGGAGACTAGGGAGTAATCCCCTTGGCGGTTAAAACGCGGGGG
ACAGCGCGTACGTGCGTTTAAGCGGTGCTAGAGCTGTCTACGACCAATTGAGCGGCCTCGGCACCGGGAT
TCTCCAGGTGTATGATGTGAACATAGCATTTATTAACAGTTCTGCTCAGTACAAATGGTAGTAGTACTAG
TGATGAGGTTACTCCGA

The sequence with high rigidity (low intrinsic cyclizability) in the 80 bp region is:

ACAGGATGTATATATCTGACACGTGCCTGGAGACTAGGGAGTAATCCCCTTGGCGGTTAAAACGCGGGGG
ACAGCGCGTACGTGCGTTTAAGCGGTGCTAGAGCTGTCTACGACCAATTGAGCGGCCTCGGCACCGGGAT
TCTCCAGGTGTATGATTCCGTGGTAAGTGATTTGAACTTTTGCTTCCTTTCGAAAATTTCAAGAATATGG
TTTGGTTGTGCTGTGCA

In both these cases, the 601 part of the sequence is denoted in black and the 80 bp segment is denoted in red. The 80 bp regions (red in the sequences above) for both these sequences were selected from among the 576 genes characterized earlier in Fig. 2b. See Extended Data Fig. 6a for plots of intrinsic cyclizability across these linker regions.

Pair 3:

The sequence with low rigidity (high intrinsic cyclizability) in the first 80 bp is:

ACAGGATGTATATATCTGACACGTGCCTGGAGACTAGGGAGTAATCCCCTTGGCGGTTAAAACGCGGGGG
ACAGCGCGTACGTGCGTTTAAGCGGTGCTAGAGCTGTCTACGACCAATTGAGCGGCCTCGGCACCGGGAT
TCTCCAGAACTTCTTTGGAGGCATCCTGAGTTGAATTAGCACTGATTGTCGTTGTATTTACAGTTTACAA
GATATAAATTATGCACT

The sequence with high rigidity (low intrinsic cyclizability) in the 80 bp region is:

ACAGGATGTATATATCTGACACGTGCCTGGAGACTAGGGAGTAATCCCCTTGGCGGTTAAAACGCGGGGG
ACAGCGCGTACGTGCGTTTAAGCGGTGCTAGAGCTGTCTACGACCAATTGAGCGGCCTCGGCACCGGGAT
TCTCCAGTTATTATTTTTTAACATTTATTGCGATGCTGCTGAAAATTTTTTTTCCACCTTGAAACTTTTC
TTTTTCCATTGAAAAAT

In both these cases, the 601 part of the sequence is denoted in black and the 80 bp segment is denoted in red. The 80 bp regions (red in the sequences above) for both these sequences were selected from among the 576 genes characterized earlier in Fig. 2b. See Extended Data Fig. 6a for plots of intrinsic cyclizability across these linker regions.

**Sliding assay:**

Each pair was considered in a separate experiment. For a given pair, the DNA fragment with high rigidity (low intrinsic cyclizability) in the 80 bp region was labeled with Cy3 at the 5' terminus distal to the 601 region, while the other member of the pair was labeled with Cy5. As a control, we swapped the dyes to rule out the possibility of a dye-dependent effect (Extended Data Fig. 7). Both fragments were mixed in equimolar amounts and nucleosomes were formed on the 601 part of the sequences via salt gradient dialysis. Sliding reactions were performed either for fixed amounts of time at various INO80 concentrations (Fig. 2f, Extended Data Fig. 7) or at saturating INO80 concentrations for various amounts of times (Extended Data Fig. 6b-c). The fact that both constructs in a pair were present simultaneously implied both were subject to exactly the same sliding conditions and for the same amounts of time, making it possible to make robust comparisons of the differences in sliding extents. See methods for the assay conditions.

**Measuring sliding extent:**

Structural and biochemical analysis suggest that INO80 moves nucleosomes in a processive, ratchet-like manner using ~10 – 20 bp step sizes, resulting in occasional intermediates between sliding start and end points[11–13].

To quantify sliding extent, a region of interest narrower than the width of the lane was selected and the median intensity across each row was plotted as a function of row number. The plot was used to subtract background and calculate peak areas. An additional faint band very close to the fully remodeled band is often visible (Fig. 2f). This could represent a near-fully remodeled intermediate. The intensities of both were added and considered as the intensity of the remodeled band ('A' In Fig. 2e-f). Likewise, a faint band is sometimes seen very close to the pre-remodeled state, which was also considered as part of the pre-remodeled configuration ('B' in Fig. 2e-f). Quantification was done on unmodified images, although the images presented in Fig. 2f, Extended Data Figs. 6, 7, and Supplementary Fig. 1 are contrast-enhanced for band visualization clarity.

**Possible reasons for reduced sliding extent in the constructs with rigid linkers:**

We discuss below several hypothetical possibilities for how rigid DNA ~40 bp ahead of a nucleosome might hinder INO80 sliding of the nucleosome towards it (Fig. 2f, Extended Data Figs. 6b-c, 7). Our data does not favor or rule out any of these possibilities.

One possibility is that rigid DNA upstream of the +1 nucleosome might reduces INO80 binding or favors dissociation. Likewise, it is also possible that an INO80 which is already sliding a nucleosome might prefer to dissociate when it engages rigid DNA ahead. However, persistence of the reduced sliding effect even under saturating INO80 conditions (Extended Data Fig. 6b-c) indicates that at least part of the effect may arise from the modulation of a reaction step that is subsequent to enzyme binding. A long pause has been detected after INO80 binding but before it has switched to a translocating mode[12], which persists for several tens of seconds even at saturating ATP. It is possible that this pause also contains a structural switch where INO80 senses and engages extranucleosomal linker DNA (via the Arp8 module). Disrupted Arp8 module:DNA binding via rigid DNA might thus disfavor formation of the translocating state. The same effect might also promote switching out of the translocating state in the case when an INO80 already sliding a nucleosome encounters rigid DNA ahead of the nucleosome (where its Arp8 module engages). Indeed pauses have also been observed in the middle of translocation events[12], which may reflect structural re-assessments of the Arp8 module:DNA interaction. It is also possible that rigid DNA directly slows down sliding speed.

Regardless of the kinetic or mechanistic details, our observations are consistent with a model where a rigid DNA segment starting ~40 bp ahead of the +1 nucleosome disfavors further upstream sliding of the nucleosome by INO80. Experiments involving, for instance, mutated INO80, much longer nucleosome constructs, or variable [ATP], could probe details of the kinetic and mechanistic pathway for this modulation, but are beyond the scope of this work.

**Supplementary Note 12: construction of the ChrV Library**

This library comprised 82,404 segments. The variable region (central 50 bp) tiles the entire length of chromosome V of *S. cerevisiae* at 7 bp resolution as follows:

Segment 1: from position 1 to 50 of chromosome V
Segment2: from position 8 to 57 of chromosome V
…
Segment 82,404: from position 576,822 to 576,871 of chromosome V

Sequences were chosen from the SacCer3 version of the genome.

The 50 bp variable regions were flanked by the standard adapters as described in supplementary note 1. See Extended Data Fig. 8 for a plot of intrinsic cyclizability as a function of position along the entire length of chromosome V in *S. cerevisiae*.

**Supplementary Note 13: plotting details for Fig. 3b**

227 genes were identified in *S. cerevisiae* chromosome V that were annotated as ORFs and had both ends mapped with high confidence[6]. Separately, the cyclizability of a fragment in the ChrV Library was assigned as the cyclizability at position n along yeast chromosome V if the fragment in question was centered around the $n^{th}$ base of chromosome V. A matrix was constructed that had 227 rows representing the 227 genes, and 1,001 columns whose values were populated with intrinsic cyclizability values from 500 bp upstream to 500 bp downstream of the dyad of the +1 nucleosome of the corresponding gene. As intrinsic cyclizabilities were measured every $7^{th}$ base, positions that had no measured cyclizability values were assigned a 'NaN' value. A 51 bp window was selected to tile the columns of this matrix. All non-NaN numbers in all rows (1 through 227) between columns 1 and 51 (including both) were considered and their mean and s.e.m calculated and used to plot the values in Fig. 3b at the x-axis location -475. Likewise, the mean and s.e.m. values plotted at x location -474 were the mean and s.e.m. of all non-NaN numbers among all rows and between columns 2 and 52 (both included), etc.

For plotting nucleosome occupancy, a similar matrix was created, except that instead of cyclizability values, it was populated with nucleosome occupancy values obtained as described in supplementary note 10.

### Supplementary Note 14: construction of Library L

The first 11,000 sequences in Library L span 500 200 bp DNA fragments that flank 500 +7 nucleosomal dyads in *S. cerevisiae*. The subsequent 44,000 sequences span these same regions, but the DNA sequences were altered by randomly selecting synonymous codons that preserve the amino acid sequences of the genes these nucleosomes lie along. We wanted to understand if alternate codons would still impart on the DNA the characteristic modulations in intrinsic cyclizability as observed in figures 3e-f.

500 +7 nucleosomes were selected at random from the genome of *S. cerevisiae*.

The DNA sequences of the set of 500 genes were altered to change codons randomly, without changing the amino acid sequences. This was done a total of 4 times, generating 4 sets of 500 codon-randomized sequences. For the first 2 sets, random codon selection accounted for the naturally occurring codon usage frequency in *S. cerevisiae*, whereas for the next two sets, all synonymous codons were selected with equal probabilities. We allowed for the possibility of selecting codons already present natively.

The coordinates of the dyads of these +7 nucleosomes were obtained from earlier works[1]. For each nucleosome, the sequence from position (dyad – 100) to (dyad + 100) was noted and oriented in the direction of transcription. This sequence (201 bp in length) was tiled as a series of 50 bp DNA fragments offset from the previous fragment by 7 bp. After covering all native sequences, the process of tiling was continued for the 4 sets of codon-altered sequences. Each of these 50 bp fragments were flanked by the standard adapters used for PCR amplification and overhang generation (supplementary note 1) and listed in the library. The following is the description of the central 50 bp region of all the library members in Library L:

Library member 1: native sequence from position 1 to 50 of the 201 bp that flank the dyad of the +7 nucleosome of the first listed transcript.

Library member 2: native sequence from position 8 to 57 of the 201 bp that flank the dyad of the +7 nucleosome of the first listed transcript

…

Library member 22: native sequence from position 148 to 197 of the 201 bp that flank the dyad of the +7 nucleosome of the first listed transcript

Library member 23: native sequence from position 1 to 50 of the 201 bp that flank the dyad of the +7 nucleosome of the second listed transcript.

…

Library member 11,000: native sequence from position 148 to 197 of the 201 bp that flank the dyad of the +7 nucleosome of the last listed transcript

Library members 11,001 – 22000: Same as from 1 to 11,000 except instead of native sequences, the first set of codon-randomized sequences were used.

Library members 22,001 – 33,000: Same as from 1 to 11,000 except instead of native sequences, the second set of codon-randomized sequences were used.

Library members 33,001 – 44,000: Same as from 1 to 11,000 except instead of native sequences, the third set of codon-randomized sequences were used.

Library members 44,001 – 55,000: Same as from 1 to 11,000 except instead of native sequences, the fourth set of codon-randomized sequences were used.

Library member 55,001 – 61,144: these sequences do not pertain to the current study.

Library members 61,145 – 67288: See Supplementary Note 8. These sequences are the same as the first 6,144 sequences in the random library (Supplementary Note 5), but with the length of the central variable region increased by half the DNA helical repeat to 55 bp.

Library member 67,289 – 83,618: these sequences do not pertain to the current study.

Library members 83,619 – 83,650: See Supplementary Note 18. These library members contain sequences that tile the 601 DNA sequence at 3 bp resolution.

Library members 83,651 – 92, 918: these sequences do not pertain to the current study.

**Supplementary Note 15: plotting details for Extended Data Fig. 10b**

In Extended Data Fig. 10b, poly dA:dT tract content, when a stretch of A or T nucleotides of length at least 4 is considered to be a poly dA:dT tract (i.e., $l$=4 condition in Extended Data Fig. 10b), was calculated as follows: for each of the 227 identified genes in *S. cerevisiae* chromosome V (supplementary note 13), the 1,001 bp DNA sequence from 500 bp upstream to 500 bp downstream of the dyad of the +1 nucleosome was noted. A row matrix containing 1,001 columns was generated whose value was 1 at locations where the second 'A' or the second 'T' in any 'AAAA' or 'TTTT' stretch occurred, and 0 at all other places. For example, if a region of DNA within one such 1,001 bp fragment was '…AACGAAAACCTTTTTG…', the corresponding entries in the row matrix would be'…0000010000011000…'. These row matrices of the 227 genes were written one below the other to construct a matrix with 227 rows and 1,001 columns. The mean poly dA:dT tract content was the column-wise mean of this matrix, smoothened over a 50 bp rolling window.

Selected genes in Extended Data Fig. 10b (red curves) that had no special poly dA:dT tract content peak in the region of the NDR were identified as follows: for each of the 227 genes, the corresponding row matrix describing the locations of AAAAs or TTTTs was noted. If the mean of the entries in this row matrix between locations 300 and 450 (i.e. from 50 bp upstream to 200 bp upstream of the +1 nucleosomal dyad) was less than or equal to 0.025, the gene was selected. In other words, selected genes represent genes where the poly dA:dT content shows no special peak in the region of the NDR. 30% of the 227 identified genes fit this criterion.

A similar process was followed when considering other values of $l$ for defining poly dA:dT tracts. For $l$ = 7 or $l$ = 10, the cutoff mean values when selecting those genes that had no special poly dA:dT tract peak in the NDR region were 0.004 and 0.002 respectively. The fraction of the 227 genes that fit this criterion were 62% and 86% respectively.

**Supplementary Note 16: comparisons of intrinsic cyclizability with predictions of earlier models and expectations**

Intrinsic cyclizability is the only mechanical property of DNA to be measured in high throughput. However, predictive models for DNA bendability have been proposed[14–17] based on common sequence features found to be associated with bent, straight or nucleosomal DNA[1,4,18–23]. Below we compare intrinsic cyclizability with such earlier known predictors of DNA mechanics.

**Periodic modulations in AT content, and overall poly dA:dT and GC content:**

Intrinsic cyclizability is independent of a particular phase imposed by the biotin tether (Extended Data Fig. 4f-g). However, it is still possible that periodic modulations in AT content, albeit at no preferred phase, influences intrinsic cyclizability. To test if this is the case, we identified the two sets of 1,000 sequences each from among the sequences in the Random Library that have the most or least values of intrinsic cyclizability. We individually plotted the AT content as a function of position along these 2,000 sequences and calculated 2,000 power spectra by taking the fast Fourier transforms of these plots. We then averaged these power spectrum plots for the two sets and found that strong and weak modulations in AT content at the helical repeat are indeed features associated with sequences that have high or low intrinsic cyclizability values respectively (Extended Data Fig. 10a). This plot is different from Extended Data Fig. 4g, where for each case, a single power spectrum is calculated from a single plot of the mean AT content as a function of position averaged over 1,000 sequences. Our finding that periodic modulations in A/T content is a feature indicative of high intrinsic cyclizability is consistent with its earlier SELEX based identification as a feature that promoted DNA looping[4].

Poly dA:dT, defined as consecutive stretches of A's or T's in a sequence, is a sequence feature present in NDRs, depleted in nucleosomes, and suggested but not directly shown to make DNA rigid[18,19,23]. We define poly dA:dT content as stretches of at least $l$ consecutive A's or at least $l$ consecutive T's, and consider various values of $l$. Higher $l$ values made the definition more restrictive. We show that poly dA:dT content is indeed high at the NDRs of chromosome V genes (Extended Data Fig. 10b). However, we find that even among the genes that have no special poly dA:dT content peak at the NDR, the NDR is still characterized by a well-defined region of low intrinsic cyclizability (Extended Data Fig. 10b). Also, while our measurements are consistent with the expectation that high poly dA:dT content makes DNA more rigid (Extended Data Fig. 10c-d), the overall poor correlation between poly dA:dT content and intrinsic cyclizability (Extended Data Fig. 10c) shows that poly dA:dT content is a poor proxy for the measured intrinsic cyclizability. Similarly, we find that the correlation between GC content and intrinsic cyclizability is also extremely poor (Extended Data Fig. 10e).

Further, as expected, we found that all gene body nucleosomes are characterized by low poly dA:dT content at the dyads. However, poly dA:dT content cannot distinguish TSS proximal from distal nucleosomes (Extended Data Fig. 10f), unlike intrinsic cyclizability (Fig. 3e). Additionally, the +1 nucleosome has the highest overall poly dA:dT which, on its own, would suggest that the physical properties of DNA disfavor the +1 nucleosome over other gene body nucleosomes. Poly dA:dT content, therefore, does not capture the details of all sequence features that could influence DNA physics, whereas intrinsic cyclizability is a directly measured quantity.

Finally, we show that codon selection for optimizing intrinsic cyclizability modulations along gene body nucleosomes (Fig. 3g) cannot be recapitulated by the optimization of known sequence features of nucleosomes such as low poly dA:dT content at dyads[19,24] and oscillatory mean AA/TT/TA contents across aligned sequences[1]. The four codon altered sets of DNA sequences spanning the 500 +7 nucleosomes continue to show low poly dA:dT content at dyads and oscillatory AA/TT/TA content (Extended Data Fig. 10g), although the intrinsic cyclizability profiles of these codon-altered sequences lack any statistically significant contrast between dyads and edges (Fig. 3g).

**<u>Dinucleotide parameters for predicting DNA curvature, flexibility and bending energy:</u>**

Dinucleotide parameters (tilt/ roll/ twist) have long been compiled and used to predict DNA curvature and flexibility in a sequence-dependent manner. We used the most recent set of reported dinucleotide parameters[25] and followed a procedure outlined recently to calculate Predicted Plectoneme Density (PPD), which is a measure of the local ease of bending DNA[26]. Briefly, local bending energy ($E_{bend}$) required to bend DNA into a loop to initiate plectoneme formation is calculated, taking into consideration DNA flexibility, and the fact that already curved regions will require less bending. These bending energies are used to assign Boltzmann-weighted probabilities $\exp\left(-\frac{E_{bend}}{k_B T}\right)$, which provides an estimate for PPD. We calculated PPD along the set of 576 genes in the Tiling Library (supplementary note 9) across which we had measured intrinsic cyclizability (Extended Data Fig. 10h). We find that PPD is a far weaker predictor of the physical properties of DNA near NDRs or at the locations of TSS-proximal nucleosomes.

**Supplementary Note 17: plotting details for Fig. 4b, 4d**

All genes in *S. cerevisiae* which had both ends mapped with high confidence[6] were considered. For every gene, all nucleosomes whose dyads lay between the TSS and the TTS were assigned a label such as +1, +2, etc depending on the order in which they occur downstream of the TSS, and also assigned to that particular TSS. Also for every ORF, all nucleosomes between the TSS and the TTS of the immediately upstream ORF, CUT or SUT (i.e., not just ORFs) were considered, irrespective of whether the upstream TTS was reported with high or low confidence[6]. These nucleosomes were labeled as -1, -2, etc, depending on the order in which they occur upstream of the TSS and also assigned to that TSS.

In figure 4b, for a particular nucleosome category as depicted on the x-axis, all nucleosomes classified under that category were first considered. From among them, those nucleosomes were chosen for which we have available (via loop-seq on the Cerevisiae Nucleosomal Library) the intrinsic cyclizability values of the 50 bp DNA fragments that lie immediately on the TSS proximal or distal side of the dyad. The mean intrinsic cyclizability values of these segments are plotted. Error bars are s.e.m.

In the first panel in figure 4d, for a particular category depicted along the x-axis, nucleosomes were first selected as described for figure 4b. From among them, only nucleosomes that 'belonged' to those TSSs whose corresponding transcripts were among the 10% most (left panel) or 3% least (middle panel) expressed among all genes in *S. cerevisiae* were selected. Expression level was defined on the basis of NetSeq measurements[7], as described in supplementary note 9.

**Supplementary Note 18: intrinsic cyclizability along the 601 DNA sequence**

We measured intrinsic cyclizability along the 147 bp 601 DNA fragment, which is known to be a strong nucleosome positioning sequence and to be highly bendable[27]. DNA on one side of the center of the 601 DNA fragment contains phased TA repeats at ~10 bp periodicity. The assumption of this TA rich side as more flexible or bendable was central to the interpretation of earlier works[28,29], and we find it to be consistent with the current analysis.

We tiled the 147 bp 601 DNA sequence as overlapping 50 bp fragments, each fragment offset from its neighbor by 3 bp, (as done in the case of the Tiling library, at 7 bp resolution (Fig. 2a)). These fragments were represented as sequence # 83,619 – 83,650 of Library L (supplementary note 14). The following is the description of the central 50 bp of sequence numbers 83,619 – 83,650 of library L:

Library member 83,619 of library L: Positions 1 – 50 of 601 DNA
Library member 83,620 of library L: Position 4 – 53 of 601 DNA
Library member 83,621 of library L: Position 7 – 56 of 601 DNA
…
Library member 83,651 of library L: Position 97 – 146 of 601 DNA

Plotted in figure 4c is intrinsic cyclizability as a function of position along 601 DNA, with a rolling window averaging over 7 fragments. The TA rich side has higher intrinsic cyclizability, consistent with earlier interpretation[28,29] that DNA on this side is more flexible or bendable.

### Supplementary Figure 1: uncropped gel images

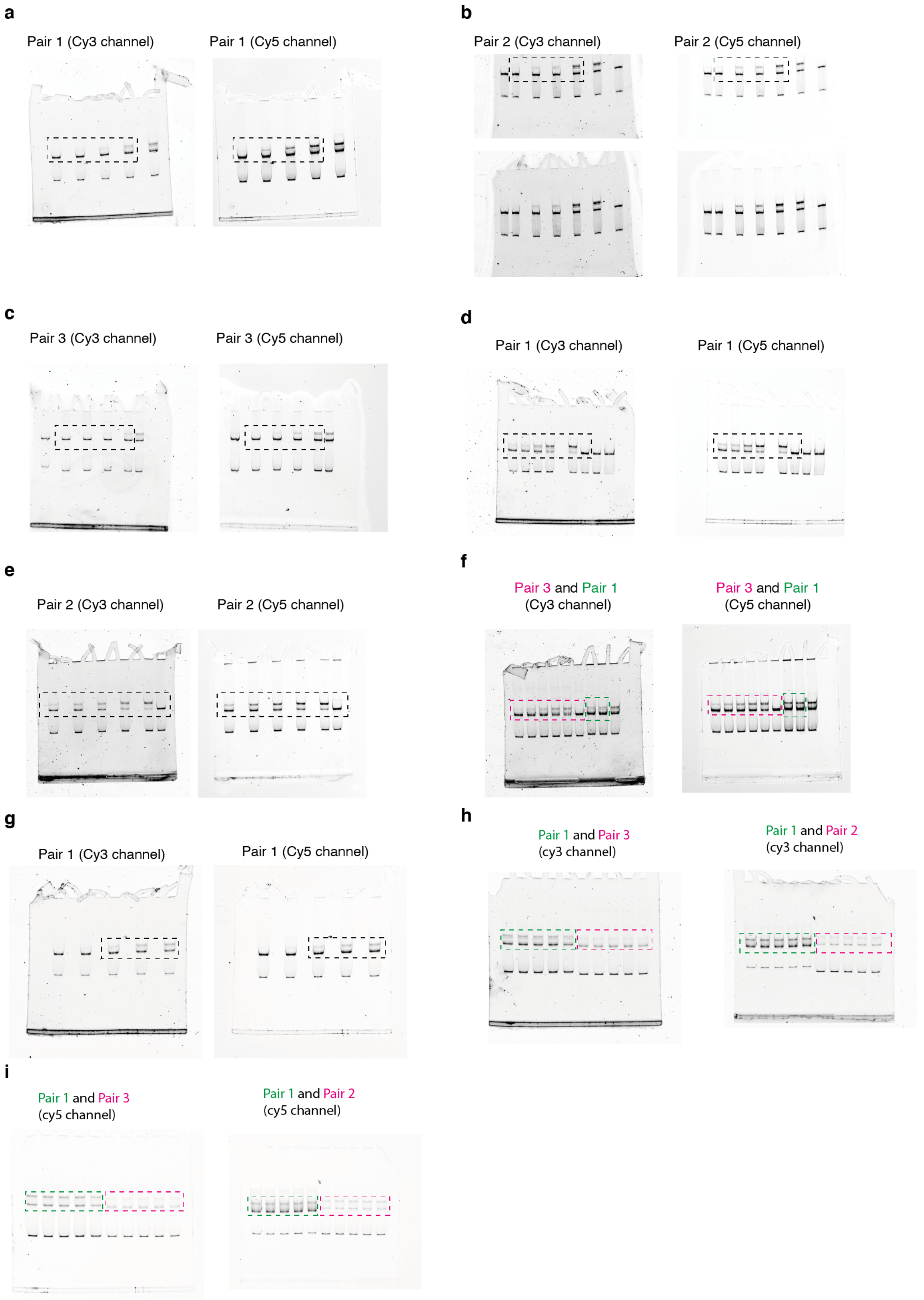

**Supplementary Fig 1**: **Uncropped gel images** (dashed rectangles indicate the cropped regions shown in Fig. 2f and Extended Data Figs. 6b and 7). **a,** The indicated cropped region is shown in Fig. 2f. Rightmost lane is for the condition [INO80] = 20 nM, but done performed side-by-side with the conditions within the dashed rectangle. **b,** Top row: The indicated cropped region is shown in Fig. 2f. [INO80] = 0.7, 20 nM in the lanes immediately to the left and right of the cropped region. The rightmost lane is a 0 ATP control. The experiments in the lanes to the right of the cropped region were not performed side by side with the other experiments. Bottom row: same as the top row, except that these images cover most of the gel. However, the top row images were the ones used for quantification purposes. **c,** The indicated cropped region is shown in Fig. 2f. The left-most lane is a nucleosome construct not part of this study. The right-most lane is for the [INO80] = 20 nM condition, not performed side-by-side with the lanes in the dashed rectangle. **d,** The cropped region is shown in Extended Data Fig. 6b. To its right is a control lacking ATP as well as INO80. The right-most lane is a nucleosome construct not part of this study. **e,** The cropped region is shown in Extended Data Fig. 6b. **f,** The cropped regions are shown (separately) in Extended Data Fig. 6b. The pink cropped region is the timecourse of INO80 sliding on nucleosomes on pair 3 at [INO80] = 30 nM. The green cropped region is for [INO80] = 30 and 60 nM. **g,** The cropped regions are shown in Extended Data Fig. 7b and refer to the dye-swapped construct version of pair 1. **h,** Pair 1, 3 in the left gel are shown in Extended Data Fig. 7a. Pair 1 in the right gel is the dye-swapped construct and shown in Extended Data Fig. 7c. Pair 2 in the right gel is shown in Extended Data Fig. 7a. **i,** Same as panel h, except that the gel is imaged for cy5 fluorescence.
